# Conserved autism-associated genes tune social feeding behavior in *C. elegans*

**DOI:** 10.1101/2023.12.05.570116

**Authors:** Mara H. Cowen, Kirthi C. Reddy, Seekanth H. Chalasani, Michael P. Hart

## Abstract

Animal foraging is an essential and evolutionarily conserved behavior that occurs in social and solitary contexts, but the underlying molecular pathways are not well defined. We discover that conserved autism-associated genes (*NRXN1(nrx-1)*, *NLGN3(nlg-1)*, *GRIA1,2,3(glr-1)*, *GRIA2(glr-2)*, and *GLRA2,GABRA3(avr-15))* regulate aggregate feeding in *C. elegans*, a simple social behavior. NRX-1 functions in chemosensory neurons (ADL and ASH) independently of its postsynaptic partner NLG-1 to regulate social feeding. Glutamate from these neurons is also crucial for aggregate feeding, acting independently of NRX-1 and NLG-1. Compared to solitary counterparts, social animals show faster presynaptic release and more presynaptic release sites in ASH neurons, with only the latter requiring *nrx-1*. Disruption of these distinct signaling components additively converts behavior from social to solitary. Aggregation induced by circuit activation is also dependent on *nrx-1*. Collectively, we find that aggregate feeding is tuned by conserved autism-associated genes through complementary synaptic mechanisms, revealing molecular principles driving social feeding.

**TEASER:** Conserved autism-associated genes mediate distinct molecular and circuit signaling components that cooperate to tune *C. elegans* social feeding behavior.

## INTRODUCTION

Social behaviors are broadly defined as interactions between individuals of the same species which can range in complexity and include mating, kin selection, parental guidance, predation, and hierarchical dominance^1,2^. One highly conserved social behavior is the formation of groups to forage or feed. Social feeding behavior is exhibited by ant colonies^3–5^, shoaling fish^6,7^, large predator herds^8–12^, and hunter-gatherer societies^13^. Social feeding can confer advantages or disadvantages depending on context, such as access to resources, predator threat, disease risk, and competition over food or mates^2,14,15^. An animal’s propensity to join a group is the result of multiple, complex, and sometimes competing environmental factors that guide their behavior^16–19^. The neuronal mechanisms underlying social feeding are not well understood, in part due to the complexity of the behavioral decisions and the underlying neuronal circuits controlling them.

The nematode *C. elegans* exhibits a wide variety of foraging behaviors and strategies^20^. For example, on a bacterial food lawn most wild isolate strains feed in large clumps of aggregating animals; however, other strains feed alone or display an intermediate level of aggregate feeding behavior^20^. Moreover, a gain-of-function polymorphism in the conserved neuropeptide receptor gene *npr-1(NPY1R)* was identified in the laboratory strain, N2 Bristol, which converts behavior from social to solitary feeding^20^. Wild social feeding behavior can therefore be genetically modeled in the solitary control strain through loss of function mutation in the *npr-1* gene (*npr-1(ad609)*)^20^. Aggregate feeding is controlled by a small sensory circuit that integrates environmental cues like oxygen levels, carbon dioxide levels, food, and aversive chemosensory stimuli, along with classical social cues like pheromones and touch^21–29^. *npr-1* modifies behavior through inhibition of RMG interneurons^21,22^ downstream of multiple highly electrically connected sensory neurons including URX, ADL, ASH, ASK, ASE, ADE, and AWB^30–32^. Moreover, the extent of social feeding is regulated by the binding affinity of *flp-21* and *flp-18* neuropeptide ligands^33^ that are released from several sensory neurons (ASE^33^, ASK^34^, ADL^35^, ASH^36^) to the *npr-1* receptor. Additionally, aggregation behavior requires the gap junction innexin gene, *unc-9,* in select neurons^30^. However, less is known about the function of chemical signaling in social feeding and how neuronal circuit properties and synapses differ between solitary and social feeders.

Individuals diagnosed with neurodevelopmental conditions, including autism, can exhibit changes in social behavior and altered sensitization to sensory stimuli^37–40^. Genomic studies have associated hundreds of genetic loci with risk for autism^41–44^, including the synaptic adhesion molecules neurexins (*NRXN1,2,3*) and their canonical post-synaptic partners, neuroligins (*NLGN1,2,3,4*)(**Supplementary Table 1**) ^45–50^. The association of neurexins and neuroligins with autism strongly suggests roles for these genes in regulating social behaviors^45–50^. Neurexins are conserved synaptic adhesion molecules that organize chemical synaptic properties including neuronal connectivity, synaptic plasticity, and excitatory/inhibitory balance^28^. Mammals have three neurexin genes that encode one long (α) and one/two short (ý and γ (specific to *NRXN1*)) isoforms of the protein^45^. Mutations in these genes in rodents alter motor activity, anxiety-like (avoidance) behavior, social approach and memory, performance of stereotyped behavior, and pre-pulse inhibition^51–58^. Neurexin mutation also impacts chemical synapse function, structure, and signaling, including presynaptic density, release probability, calcium dynamics, and post-synaptic currents^55–62^.

*C. elegans* has a single ortholog of neurexins, *nrx-1*, which is 27% identical to human NRXN1 at the amino acid level based on DIOPT alignment^63^, with nearly identical domain structure (**Supplementary Table 1**)^64^. In *C. elegans, nrx-1* contributes to retrograde inhibition of neurotransmitter release at neuromuscular junctions, regulation of GABA receptor diffusion and alignment of GABA, synaptic clustering, and synapse formation^65–70^. However, these synaptic functions of *nrx-1* have rarely been linked to distinct behaviors, with the exception of male mating, where *nrx-1* impacts male response to hermaphrodite contact^71^ and time to spicule protraction^72^. Despite these advances, we still have much to learn about the functions of *nrx-1* in circuits and synapses outside of the neuromuscular junction and how *nrx-1* mechanistically alters complex behaviors.

Using *npr-1(ad609)* mutant *C. elegans* to model social feeding behavior, we find three molecularly independent synaptic mechanisms (synaptic adhesion molecules NRX-1 and NLG-1 and the classical excitatory neurotransmitter, glutamate) that work together to tune foraging behavior from solitary to social. Through mechanistic study, we also identify the downstream glutamate receptors that regulate aggregation behavior, homologs of which are also associated with autism, highlighting conservation of this pathway across species. Despite *nrx-1* and the vesicular glutamate transporter required for glutamate release, *eat-4*, both functioning in ASH and ADL sensory neurons to modulate aggregation behavior, the mechanisms by which they control social feeding are distinct - faster glutamate release dynamics occur independently of *nrx-1* while higher number of pre-synaptic release sites depends on *nrx-1*. These additive neuronal mechanisms exemplify the complexity of *C. elegans* foraging strategies and provide insights into how variation in social behavior is achieved at the genetic, molecular, and circuit levels.

## RESULTS

### NRX-1(α) functions in ADL and ASH sensory neurons for aggregation behavior

Neurexin genes, including *nrx-1* in *C. elegans,* are broadly expressed in neurons in mammals and invertebrates. We used a database of neuronal gene expression profiles for every neuron (CENGEN)^73^ to confirm that *nrx-1* transcripts are present in the RMG interneurons and upstream sensory neurons implicated in aggregation behavior (**Fig. 1A**). Given the broad expression of *nrx-1* in RMG interneurons and its synaptic partners, we asked if *nrx-1* functions in aggregation behavior. We quantified aggregation behavior as the number of *C. elegans* in contact with two or more animals, based on previous literature^20^, for 50 day 1 adults, using longitudinal, blinded image analysis (**Fig. 1B**). As expected, we find that *npr-1(ad609)* mutants aggregate significantly more than solitary controls (*npr-1* average = 38.67, SEM= 1.675 vs. solitary control average =2.2, SEM =0.860 (**Fig. 1D&E**). Solitary controls consist of the N2 Bristol strain and solitary animals from the N2 background with an integrated transgene (*otIs525*) and/or *him-8* mutation, used for genetic crosses. Aggregation behavior was not impacted by *otIs525;him-8* in the solitary (N2) or aggregating background (*npr-1(ad609)*)(**Supplemental Fig. 1A**).We tested three mutant alleles of *nrx-1*: a large deletion in *nrx-1* that disrupts both the long (α) and short (γ) isoforms *(wy778)*, an α-isoform specific deletion *(nu485),* and a nonsense mutation leading to a premature stop codon early in the α-isoform *(gk246237)*(**Fig. 1C**). In the *npr-1(ad609)* aggregating background, all three alleles of *nrx-1* significantly decreased the number of aggregating *C. elegans* compared with *npr-1(ad609)* alone (**Fig. 1D&E**). Notably, *npr-1(ad609)* animals carrying any of the *nrx-1* mutant alleles show intermediate aggregation behavior compared to solitary controls or *nrx-1* mutants alone, which show almost no aggregation behavior. Thus, we find that *nrx-1* is essential for aggregation behavior induced by *npr-1* mutation, such that disruption of *nrx-1* reduces aggregation behavior of *npr-1* animals by ∼40%, which is primarily mediated by the α isoform. Animals carrying the *npr-1* variant of a wild social isolate strain (215F in Hawaiian CB4856) in an otherwise N2 background (*qgIR1*) also display aggregation behavior (**Supplemental Fig. 1B&C**). Aggregate feeding in this strain is dependent on *nrx-1*, confirming that *nrx-1* contributes to social feeding.

**Fig. 1.**
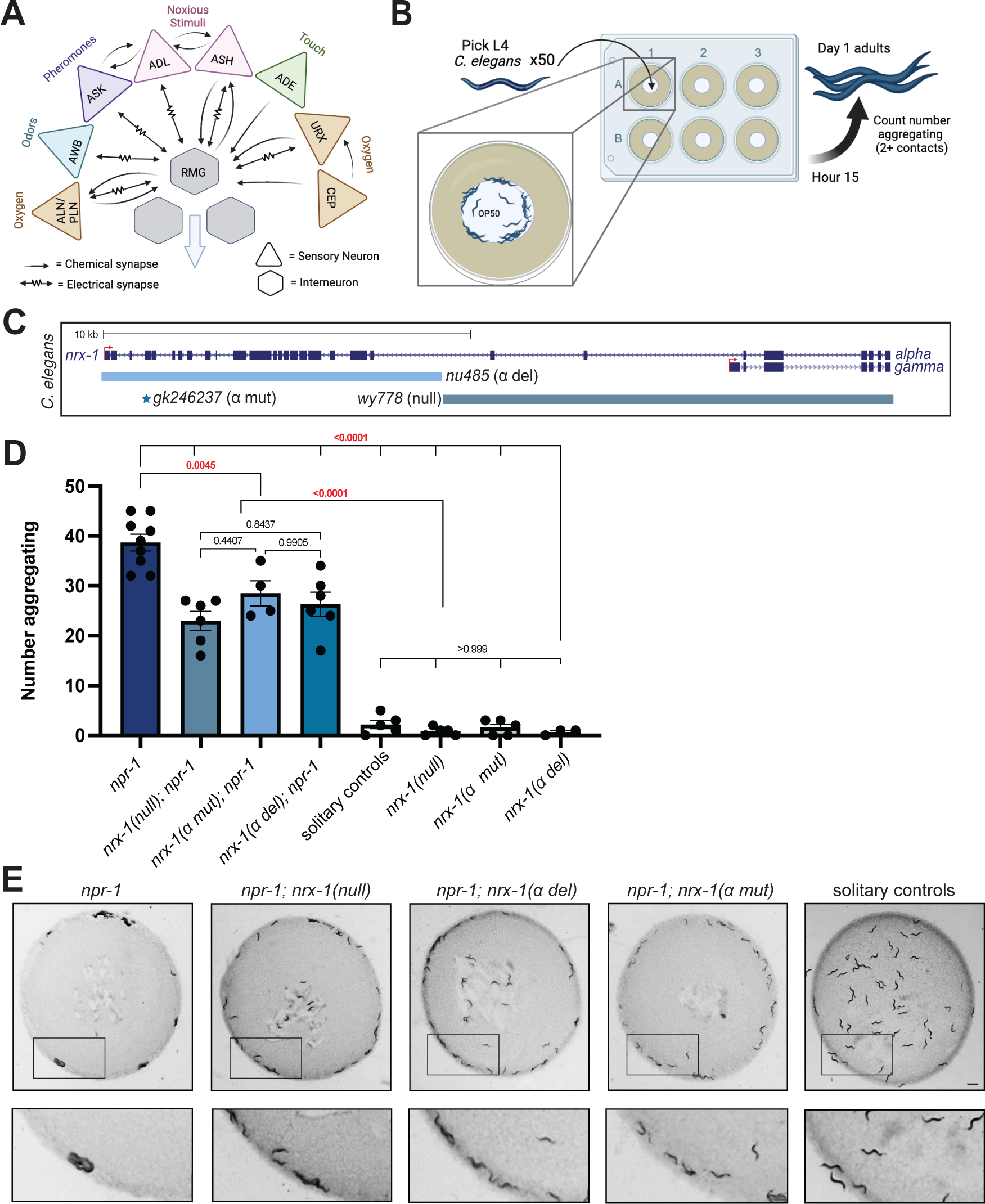
*nrx-1* is essential for aggregation behavior. **A)** Circuit diagram of sensory integration circuit. Connectome based on NemaNode and WormWiring data. **B)** Cartoon of medium through-put aggregation behavior assay with 50 timed day 1 adult worms per well of a 6-well WormCamp then imaged using WormWatcher platforms and scored for aggregation behavior defined as two or more animals in direct contact. **C)** Schematic of *C. elegans nrx-1* gene showing mutant alleles used and isoforms removed by functional null and α-isoform specific mutants. **D)** Graph showing number of aggregating animals in various genetic backgrounds. All mutant *nrx-1* alleles (*wy778 = nrx-1 null, gk246237 = nrx-1* α *mut*, *nu485 = nrx-1* α *del*) show decreased aggregation behavior. **E)** Representative images of aggregation behavior in *npr-1(ad609), npr-1(ad609);nrx-1(wy778), npr-1(ad609);nrx-1(nu485)*, *npr-1(ad609);nrx-1(gk246237)* mutants, and solitary controls (Scale bar = 1mm).

To localize the function of *nrx-1* in aggregation behavior, we created animals expressing *nrx-1* isoforms under various neuron-specific promoters. We tested a large panel of promoters and quantified the impact on aggregation behavior of *nrx-1* nulls in the *npr-1* aggregating background (**Supplemental Fig. 1D**). Expression of NRX-1(α) in all neurons using the *ric-19* promoter completely restored aggregation behavior in *npr-1*; *nrx-1(wy778)* mutants to the level of aggregating *npr-1(ad609)* animals (**Fig. 2A-C**). Expression of NRX-1(γ) in neurons under the same *ric-19* promoter had no impact, confirming a specific role for the α-isoform in aggregation behavior (**Fig. 2A&B**). Further, expressing the α-isoform of NRX-1 in the RMG interneurons and several sensory neurons including ADL and ASH (*flp-21p*), or in both ADL and ASH sensory neurons (*nhr-79p*), restored aggregation behavior to levels comparable to pan-neuronal expression (**Fig. 2A-C**). Collectively, these data suggest that NRX-1(α) functions in at least in two pairs of sensory neurons for aggregation behavior.

**Fig. 2.**
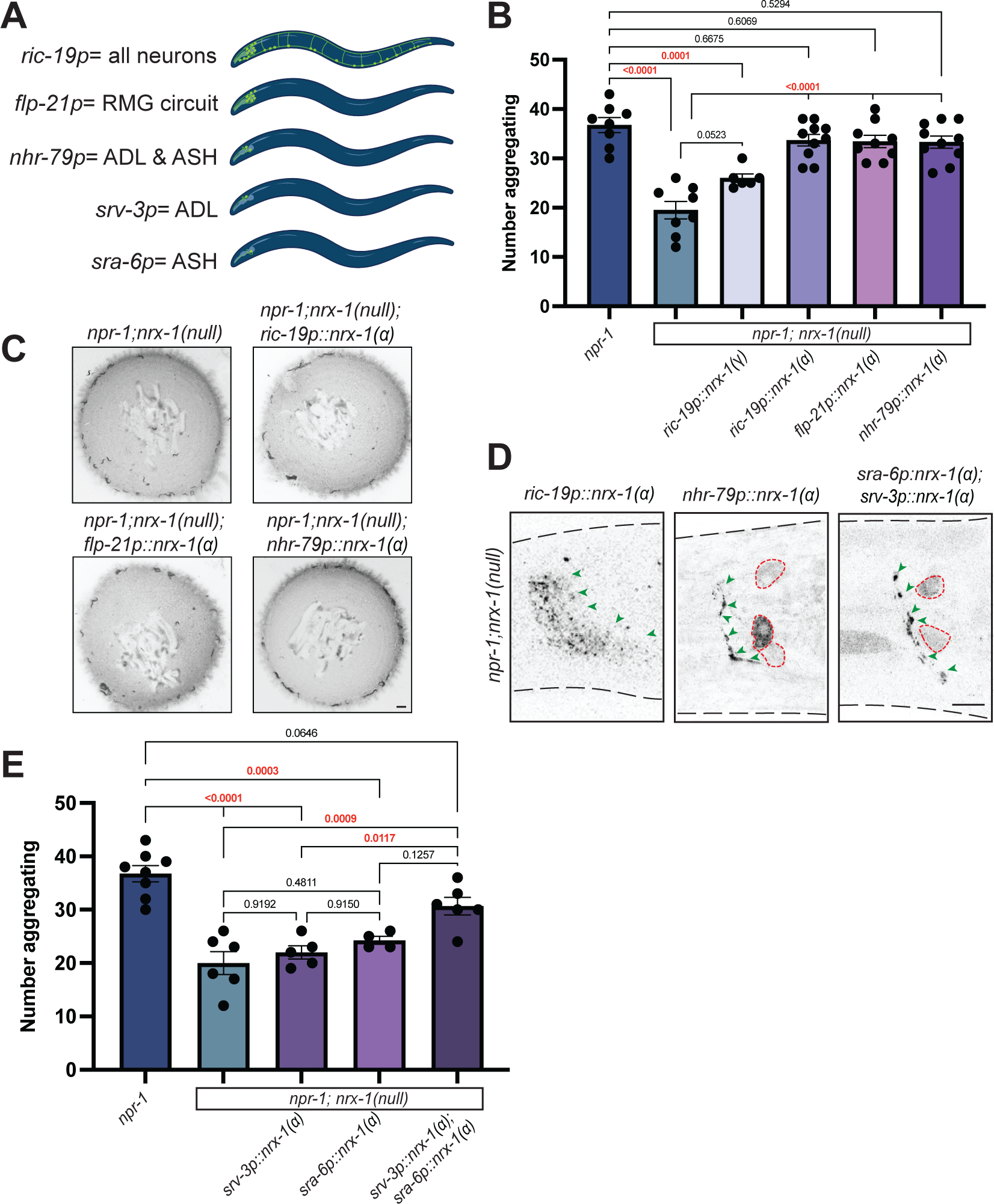
NRX-1(α) acts in ADL and ASH sensory neurons for aggregate feeding. **A)** Cartoons showing the neurons where each promoter is expressed. *ric-19p* expresses in all neurons, *flp-21p* expresses in neurons in several sensory neurons and RMG, *nhr-79p* expresses in ADL and ASH sensory neurons, *srv-3p* expresses in ADL neurons, and *sra-6p* expresses in ASH neurons. Graph showing number of aggregating animals **(B)** and representative images of aggregation behavior assay plates **(C)** in *npr-1(ad609);nrx-1(null*) mutants with NRX-1(α) driven by *ric-19, flp-21,* and *nhr-79* promoters, and NRX-1(γ) driven by the *ric-19* promoter and controls (Scale bar = 1mm). **D)** Confocal image of NRX-1(α) expression in all neurons (*ric-19p::sfGFP::nrx-1*), ADL and ASH neurons (*nhr-79p::sfGFP::nrx-1),* and ADL and ASH neurons (*sra-6p::sfGFP::nrx-1 & srv-3p::sfGFP::nrx-1)*. Green arrows indicate NRX-1 axonal expression. Red dashed lines show cell bodies. *ric-19p::sfGFP::nrx-1(α)* imaging performed *in nrx-1(wy778)* (Scale bar = 10μm). **E)** Graph showing number of aggregating animals in various genetic backgrounds. Data for *npr-1* and *npr-1;nrx-1* is plotted in both 2B and 2E.

We confirmed expression and localization of the various *nrx-1* transgenes by fusing a Superfolder GFP to the *nrx-1* coding sequence and monitoring fluorescence in the various neurons (**Fig. 2D, Supplemental Fig. 1E**)^74^. In all transgenic animals sfGFP::NRX-1(α) localized along the neurites and processes of the neurons in a punctate pattern; with some expression also observed within the cell body (**Fig. 2D**). To determine if *nrx-1* functions in ADL and/or ASH neurons for aggregation behavior, we expressed sfGFP::NRX-1(α) specifically in ADL using the *srv-3* promoter or specifically in ASH using the *sra-6* promoter. Expression of sfGFP::NRX-1(α) in ADL or ASH individually did not restore aggregation behavior to *npr-1* levels, however, combination of these same two transgenes increased aggregation behavior, confirming the function of NRX-1(α) in both pairs of sensory neurons (**Fig. 2D-E**). These data are consistent with previous results showing that ablating both ADL and ASH disrupts aggregation behavior^28^.

### NLG-1 is essential for aggregation behavior independently of NRX-1

*nlg-1* is the single *C. elegans* ortholog of the neuroligin synaptic adhesion genes *NLGN1,2,3,4*^75^, the well-characterized trans-synaptic partner of *NRXN1*(*nrx-1)*^76^. Using a large deletion in *nlg-1*(*ok259*)^75^, we asked if disruption of *nlg-1* also alters aggregation behavior in *npr-1(ad609)* mutants. We find that loss of *nlg-1* leads to a significant decrease in aggregation behavior of *npr-1(ad609)* mutant animals but has no effect in solitary control animals (**Fig. 3A,B,&E**). To localize the function of *nlg-1* in aggregation behavior we used a similar transgenic rescue approach as for *nrx-1*. Expression of sfGFP::NLG-1 in all neurons using the *ric-19* promoter partially restored aggregation behavior (**Fig. 3C**). Expression of sfGFP::NLG-1 in ADL and ASH sensory neurons (*nhr-79* promoter) or in the RMG interneurons (*nlp-56* promoter) did not impact aggregation behavior (**Fig. 3C**). Expressing sfGFP::NLG-1 in ADL (*srv-3* promoter) or ASH (*sra-6* promoter) individually or in AIA (*ins-1* promoter) did not rescue aggregation behavior (**Supplemental Fig. 2A**). We also confirmed expression of all sfGFP::NLG-1 transgenes by analyzing expression by the sfGFP tag (**Supplemental Fig. 2B**). Together, these results imply that NLG-1 in neurons is sufficient to partially modify aggregation behavior.

**Fig.3.**
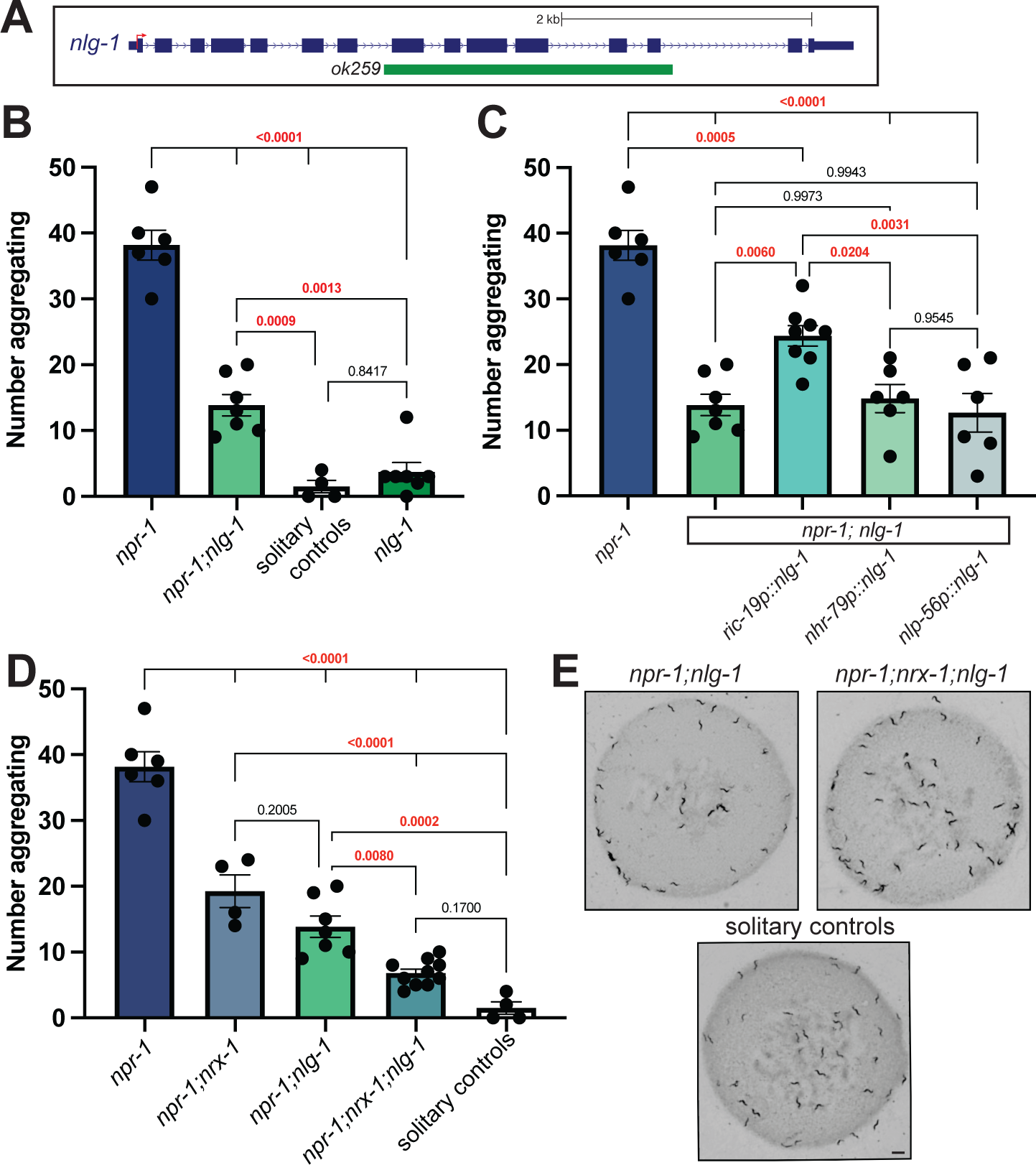
NLG-1 contributes independent of NRX-1 in aggregation behavior. **A)** Schematic of *C. elegans nlg-1* gene showing deletion allele assessed. **B)** Graph showing number of aggregating animals in *npr-1(ad609)*, *npr-1(ad609);nlg-1(ok259)*, *nlg-1(ok529)*, and solitary controls. *nlg-1* deletion decreased aggregation behavior in *npr-1* animals. **C)** Graph showing number of aggregating animals in *npr-1(ad609);nlg-1(ok259)* mutants with NLG-1 driven by *ric-19, nhr-79,* and *nlp-56* promoters and controls. *ric-19p* expresses in all neurons, *nhr-79p* expresses in ADL and ASH sensory neurons and *nlp-56p* expresses in RMG neurons. **D)** Graph showing number of aggregating animals in *npr-1(ad609)*, *npr-1(ad609);nrx-1(wy778)*, *npr-1(ad609);nlg-1(ok259), npr-1(ad609);nrx-1(wy778);nlg-1(ok259)*, and solitary controls. **E)** Representative images of aggregation behavior in *npr-1(ad609);nlg-1(ok259), npr-1(ad609);nrx-1(wy778); nlg-1(ok259)* and solitary controls (Scale bar = 1mm). Data for *npr-1* and *npr-1;nlg-1* is plotted in 3B, 3C, and 3D. Data for solitary controls is plotted in 3B and 3C.

To test whether *nrx-1* and *nlg-1* function together, we created an *npr-1(ad609);nrx-1(wy778);nlg-1(ok259)* triple mutant. We find a significant decrease in aggregation behavior in the triple mutant animals compared to either double mutant (**Fig. 3D&E**). These findings suggest that both *nrx-1* and *nlg-1* are critical for aggregation behavior, but likely function in parallel, non-epistatic, molecular pathways. These data are surprising, but not inconsistent with previous studies finding that *nrx-1* and *nlg-1* can function together^70,77^, independently^67–69^, or even antagonistically^71,72^.

### Glutamate signaling from ADL and ASH neurons is necessary for aggregation behavior

ADL and ASH sensory neurons signal via glutamate to modify animal behavior and silencing the gap junctions in ADL and ASH has been shown to not impact social feeding behavior^30^, likely implicating chemical signaling from these neurons. We hypothesized that mutations in the glutamate transporter EAT-4, a homolog of the human VGLUTs, might also affect aggregation behavior. Disrupting *VGLUT*(*eat-4)* in an aggregating *npr-1* background significantly decreases aggregation behavior compared to *npr-1* mutants (**Fig. 4A&C**). To test if glutamate functions specifically in ADL and ASH for aggregation behavior, we expressed EAT-4 using the *nhr-79* promoter which restored aggregation behavior of *npr-1(ad609); eat-4(ky5)* double mutants to the same level as *npr-1* mutants (**Fig. 4A&C**). Like NRX-1, we find that expression of EAT-4 is needed in both ADL and ASH neurons, where expression in either neuron alone is insufficient to restore aggregation behavior of *npr-1(ad609); eat-4(ky5)* mutants (**Fig. 4A**). We confirmed expression of all EAT-4 transgenes with visualization of a trans-spliced GFP (**Supplemental Fig. 3**).

**Fig. 4.**
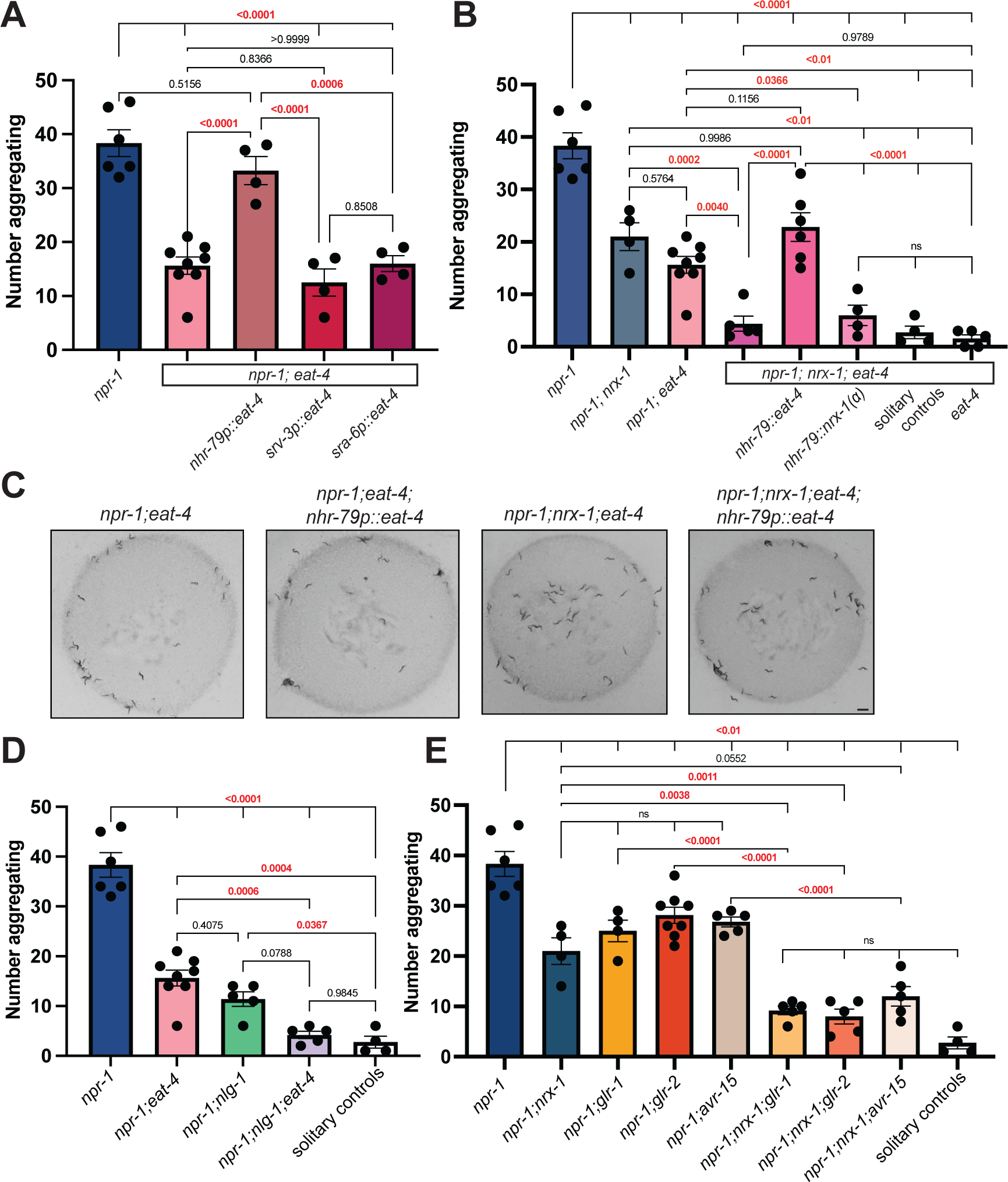
Aggregation behavior depends on glutamate signaling from ADL and ASH neurons. **A)** Graph showing number of aggregating animals in *npr-1(ad609)* compared to *npr-1;eat-4(ky5)* mutants and number of aggregating animals in *npr-1(ad609);eat-4(ky5)* mutants with EAT-4 driven by *nhr-79, srv-3,* and *sra-6* promoters. **B)** Graph showing number of aggregating animals in *npr-1(ad609)*, *npr-1(ad609);nrx-1(wy778)*, *npr-1(ad609);eat-4(ky5)*, *npr-1(ad609);nrx-1(wy778);eat-4(ky5)* mutants. Graph also includes *npr-1(ad609);nrx-1(wy778);eat-4(ky5)* mutants with EAT-4 driven under the *nhr-79* promoter, *npr-1(ad609);nrx-1(wy778);eat-4(ky5)* mutants with NRX-1(α) driven under the *nhr-79* promoter, and solitary controls. **C)** Representative images of aggregation behavior in *npr-1(ad609);eat-4(ky5)*, *npr-1(ad609);eat-4(ky5*); *nhr-79p::eat-4*, *npr-1(ad609);nrx-1(wy778);eat-4(ky5),* and *npr-1(ad609);nrx-1(wy778);eat-4(ky5); nhr-79p::eat-4* animals (Scale bar = 1mm). **D)** Graph showing number of aggregating worms in *npr-1(ad609), npr-1(ad609);eat-4(ky5), npr-1(ad609);nlg-1(ok259), npr-1(ad609);nlg-1(ok259);eat-4(ky5)* mutants, and solitary controls. **E)** Graph showing number of aggregating animals in *npr-1(ad609), npr-1(ad609);nrx-1(wy778), npr-1(ad609);glr-1(n2461), npr-1(ad609);glr-2(ok2342), npr-1(ad609);avr-15(ad1051), npr-1(ad609);nrx-1(wy778);glr-1(n2461), npr-1(ad609);nrx-1(wy778); glr-2(ok2342)*, and *npr-1(ad609);nrx-1(wy778);avr-15(ad1051)* mutants. Data for *npr-1* and *npr-1;eat-4* is plotted in 4A, 4B, and 4D. Data for *npr-1;nrx-1* is plotted in 4B and 4E. Data for solitary controls is plotted in 4B, 4D, and 4E.

The shared role of *nrx-1* and *eat-4* in ADL and ASH sensory neurons suggested that *nrx-1* and *eat-4* may function together in these neurons to regulate aggregation behavior. However, we find that *npr-1(ad609)*; *nrx-1(wy778)*; *eat-4(ky5)* triple mutants further reduce aggregation behavior compared to either *npr-1(ad609); nrx-1(wy778)* or *npr-1(ad609); eat-4(ky5)* double mutants (**Fig. 4B**). This result indicates that glutamate and *nrx-1* may function in parallel, non-epistatic, pathways to affect aggregation behavior. Since we find that *nlg-1* and *nrx-1* also function independently, we asked if *eat-4* and *nlg-1* may function through the same molecular pathway. However, we find that perturbing glutamate signaling and *nlg-1* in an *npr-1(ad609);eat-4(ky5);nlg-1(ok259)* triple mutant further decreases aggregation behavior, to a level similar to that observed in solitary controls (**Fig. 4D**). Therefore, we conclude that there are three novel molecular signaling components that contribute to aggregation behavior and find that *nrx-1*, *nlg-1*, and *eat-4* function in genetically distinct or parallel pathways. Remarkably, we find that loss of each component individually reduces aggregation behavior significantly, but combination of any two reduces aggregation behavior further towards solitary behavior. This demonstrates that aggregation behavior is regulated by heterogenous genetic pathways which together tune behavior between solitary and social feeding.

To further explore the interplay of *nrx-1* and glutamate in ADL and ASH sensory neurons, we expressed EAT-4 or NRX-1(α) in these neurons of *npr-1(ad609)*; *nrx-1(wy778)*; *eat-4(ky5)* triple mutants using the *nhr-79* promoter. Expression of EAT-4 in ADL and ASH in the *npr-1(ad609)*; *nrx-1(wy778)*; *eat-4(ky5)* triple mutants restored aggregation behavior to the level of *npr-1(ad609);nrx-1(ad609)* (**Fig. 4B**), providing further evidence that the role of glutamate in aggregation behavior is independent of *nrx-1* despite functioning in the same sensory neurons. Expression of NRX-1(α) in ADL and ASH in *npr-1(ad609)*; *nrx-1(wy778)*; *eat-4(ky5)* triple mutants did not alter aggregation behavior (**Fig. 4B**), suggesting a possible dependence of *nrx-1* on glutamate. Together with the additive behavioral findings for *nrx-1* and *eat-4,* this result implies a dual role for *nrx-1* in aggregation behavior − one dependent on glutamate and one independent of glutamate that may occur in non-glutamatergic neurons.

### Multiple glutamate receptors regulate aggregation behavior

Our results thus far have focused on the pre-synaptic mechanisms regulating aggregation behavior. To explore how aggregate feeding is controlled on the post-synaptic side, we next tested the role of glutamate receptors. We analyzed mutants in glutamate receptors including *GRIA1,2,3(glr-1), GRIA2(glr-2), GRIN2B(nmr-2), GRM3(mgl-1)*, and *GLRA2,GABRA3(avr-15)*.

We find that *glr-1(n2461), glr-2(ok2342),* and *avr-15(ad1051),* but not *mgl-1(tm1811)* or *nmr-2(ok3324)* reduce aggregation behavior in the *npr-1(ad609)* background (**Fig. 4E**, **Supplemental Fig. 3B**). Notably, while *glr-1* and *glr-2* are excitatory AMPA-like receptors^78^, *avr-15* is an inhibitory glutamate-gated chloride channel^79^ suggesting that a complex balance of glutamate signaling is involved in aggregation behavior.

We next wondered whether *nrx-1* may function at the level of post-synaptic glutamate receptor function or clustering similar to its role at other synapses^80,81^. To answer this, we created triple mutants for *npr-1, nrx-1*, and each glutamate receptor. We find *nrx-1(wy778)* with each glutamate receptor mutation further reduces aggregation behavior compared with *nrx-1* or each respective receptor mutant alone in an aggregating background (**Fig. 4E**). These data suggest that *nrx-1* acts additively with the receptors, where loss of a single receptor reduces aggregation behavior, and loss of *nrx-1* may lower functionality of the other two remaining receptors or act through independent mechanisms as indicated by the results with loss of glutamate itself.

### Glutamate release is higher in aggregating *C. elegans*

To determine how glutamate signaling contributes to solitary versus aggregate feeding behavior, we used fluorescence recovery after photobleaching (FRAP) of the pH-sensitive GFP-tagged vesicular glutamate transporter, EAT-4::pHluorin^82^. To gain temporal information of synaptic release, we photobleached fluorescence at ASH pre-synaptic sites and recorded fluorescence recovery for two minutes post bleach (**Fig. 5 A&B**). Recovery was normalized to pre-bleach fluorescence as the maximum (1) and post-bleach fluorescence as the minimum (0)^83^. The slope of the recovery allowed us to compare rates of ASH glutamate release between genotypes. Initial EAT-4::pHluorin levels in ASH were not different between genotypes (**Fig. 5C**). We find that ASH neurons have faster spontaneous glutamate release in aggregating *npr-1(ad609)* animals compared to solitary controls as exemplified by greater overall and faster fluorescence recovery (**Fig. 5D**). We next asked whether NRX-1 had a role in the increased rate of glutamate release and find that ASH neurons in *npr-1(ad609);nrx-1(wy778)* mutants also have faster glutamate release dynamics relative to solitary controls (**Fig. 5E**). *nrx-1(wy778)* mutants in a solitary background had similar ASH glutamate release dynamics to that of solitary controls (**Fig. 5D**). Notably, we find that glutamate release is higher in strains generated in an aggregating background (*npr-1(ad609)* or *npr-1(ad609);nrx-1(wy778)*) strains compared to strains generated in the solitary background (N2 and *nrx-1(wy778)*). Therefore, while aggregation behavior is affected by *nrx-1*, glutamate dynamics occur independent of *nrx-1,* providing further evidence that *nrx-1* and glutamatergic signaling regulate aggregate feeding through distinct mechanisms.

**Fig. 5.**
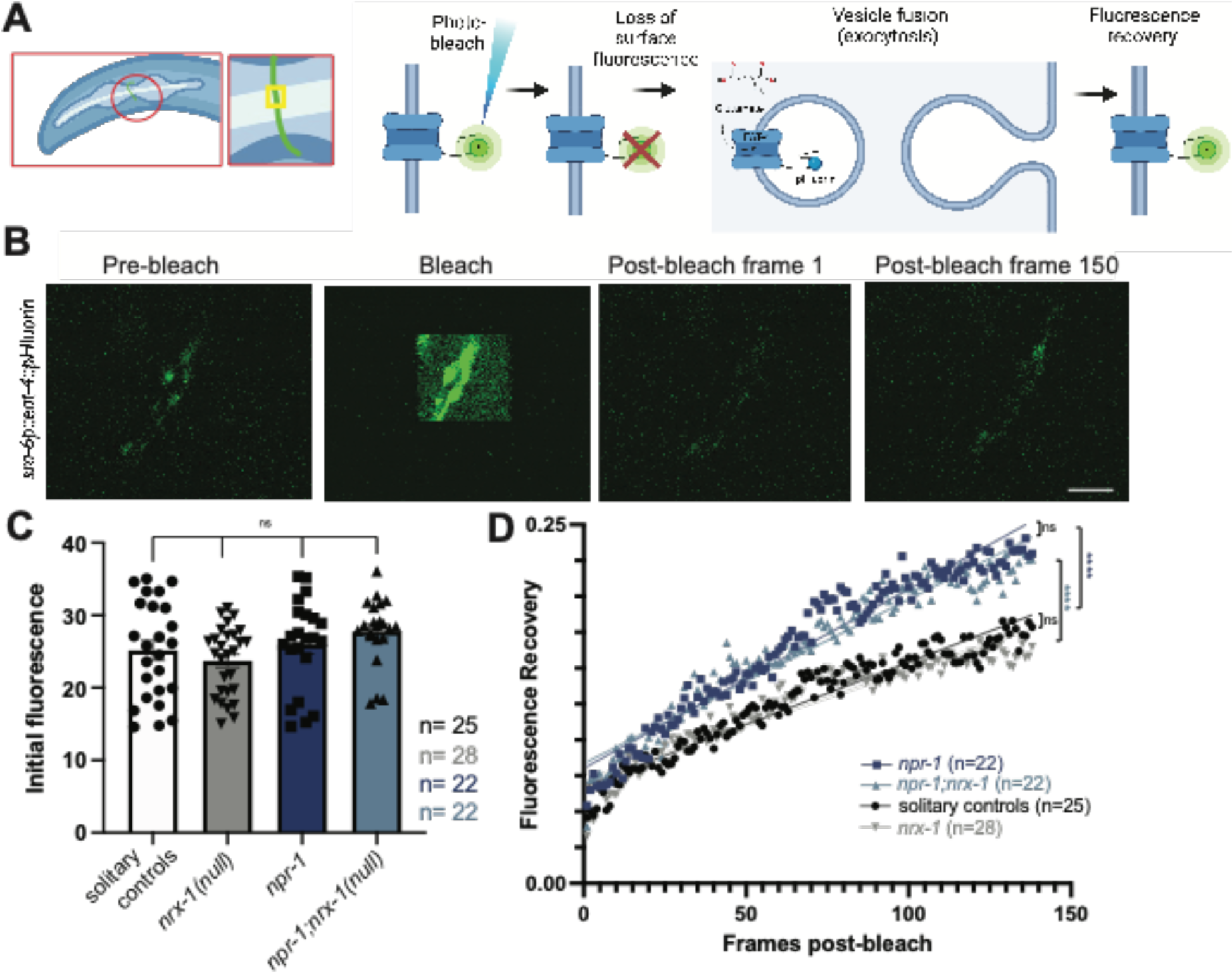
Glutamate release is faster in aggregating *C. elegans,* independent of NRX-1. **A)** Cartoon of *sra-6p::eat-4::pHluorin* experiment, including schematic of small neuron section bleached and EAT-4::pHluorin photobleaching and recovery process. **B)** Representative images of ASH neuron prior to bleaching (pre-bleach), during bleach, immediately following bleach, and after recovery period of two minutes (Scale bar = 5μm). **C)** Graph showing initial fluorescence values taken from first 10 pre-bleach frames of FRAP experiments. **D)** Graph of post-bleach recovery as a fraction of initial fluorescence by post-bleach frame up to frame 138 (120 seconds, frame taken every 0.87 seconds) Comparisons shown on graph include: *npr-1* and *npr-1;nrx-1* (p=0.278), solitary control and *nrx-1*(p=0.080), solitary control and *npr-1* (dark blue, p<0.0001), and *npr-1;nrx-1* and *nrx-1* (light blue, p<0.0001).

### ASH pre-synaptic puncta are higher in aggregating *C. elegans* dependent on NRX-1

To investigate whether *nrx-1* alters aggregation behavior through a role in synaptic structure or architecture, we analyzed pre-synaptic morphology of the ADL and ASH neurons using enhanced resolution confocal microscopy of the chemical GFP-tagged pre-synaptic marker clarinet CLA-1 (a bassoon ortholog)(**Fig. 6A**)^84^. Specifically, we quantified CLA-1::GFP puncta in the neurites of ADL or ASH sensory neurons using the *srv-3* and *sra-6* promoters respectively, via an unbiased particle analysis (see methods for details, **Fig. 6A**). We find no significant difference in ADL pre-synaptic puncta number between aggregating, solitary, or *nrx-1* mutants (**Fig. 6B&C**). We next quantified pre-synaptic puncta in ASH neurons, and unlike ADL, we find that aggregating *npr-1(ad609)* mutants have a significant increase in the number of CLA-1::GFP puncta compared with solitary controls (**Fig. 6D&E**). Further, the number of ASH CLA-1::GFP puncta in a *npr-1(ad609);nrx-1(wy778)* double mutant was significantly lower than in *npr-1* alone (**Fig. 6D&E**). These results indicate that aggregating animals have more ASH pre-synaptic puncta than solitary controls and that this increase is dependent on NRX-1. The impact of *nrx-1(wy778)* on CLA-1::GFP puncta in ASH was also context dependent and only altered puncta number in the aggregating strain with no impact in the solitary control background.

**Fig. 6.**
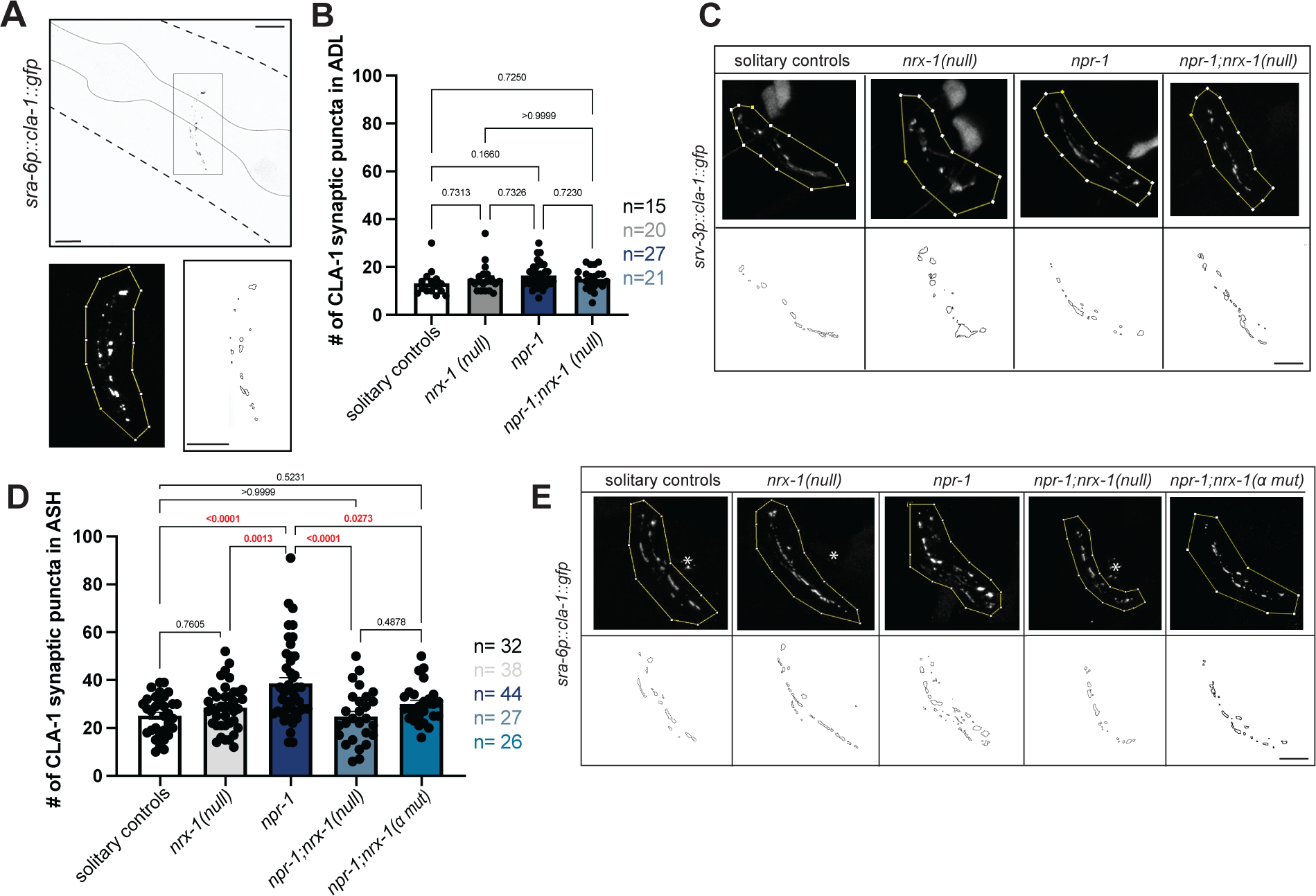
Higher number of ASH pre-synaptic puncta in aggregating *C. elegans* depends on *nrx-1*. **A)** Confocal micrograph of *sra-6p::cla-1::gfp* construct with pharynx outlined. Region of interest (ROI) in which counts are performed and puncta outlines generated by FIJI. Soma and projections outside of the nerve ring are not included in ROI (Scale bar = 10μm). Graph showing number **(B)** and representative images **(C)** of *srv-3p::cla-1::gfp* puncta in ADL in solitary controls *nrx-1(wy778), npr-1(ad609)*, and *npr-1(ad609);nrx-1(wy778)* mutants. Graph showing number **(D)** and representative images **(E)** of *sra-6p::cla-1::gfp* puncta in ASH in solitary controls *nrx-1(wy778), npr-1(ad609)*, and *npr-1(ad609);nrx-1(wy778)* mutants (Scale bars = 10μm).

To determine if a specific isoform of NRX-1 is responsible for regulating the higher number of pre-synaptic puncta number in aggregating strains, we tested an α-isoform specific mutant allele, *nrx-1(gk24623)*. We find that *npr-1(ad609);nrx-1(gk24623)* mutants had fewer ASH CLA-1::GFP puncta relative to *npr-1(ad609)* aggregating animals, similar to what we observed in *npr-1(ad609)* animals carrying the null allele of *nrx-1* that knocks out both α and γ isoforms (**Fig. 6D&E**). This result suggests that, like aggregation behavior, pre-synaptic architecture, which is modified in aggregating compared to solitary strains, is selectively mediated by NRX-1(α).

### Aggregation behavior induced by activation of sensory neurons and RMG interneurons depends on NRX-1

Increased glutamate release dynamics and pre-synaptic puncta in *npr-1* animals likely promote neuronal signaling from ASH to other neurons (i.e. ADL, RMG) in the sensory integration circuit that regulates aggregation behavior. Previous work found that activating sensory neurons and the RMG interneurons through expression of a constitutively active Protein Kinase C *(flp-21p::pkc-1(gf)),* induces aggregation behavior in solitary animals by increasing release of neurotransmitters and neuropeptides^21^. We queried if *nrx-1* was needed for aggregation behavior outside of the context of *npr-1* and find that, as previously reported, *flp-21p::pkc-1(gf)* induces social feeding, albeit at lower levels than *npr-1(ad609)* mutants^21^ **(Fig. 7A&B**). Lastly, we show that *nrx-1(wy778); flp-21p::pkc-1(gf)* animals aggregate less than *flp-21p::pkc-1(gf)* alone **(Fig. 7 A&B)**. Therefore, *nrx-1* is necessary for aggregation behavior induced by increased neuronal signaling within the sensory integration circuit that drives aggregation behavior. These results complement our finding that aggregating animals shift their behavior towards a solitary state when the number of pre-synaptic release sites is decreased in *nrx-1* mutants, by showing that *nrx-1* mutations prevent the conversion of solitary to more social behavior when circuit activity is increased.

**Fig. 7:**
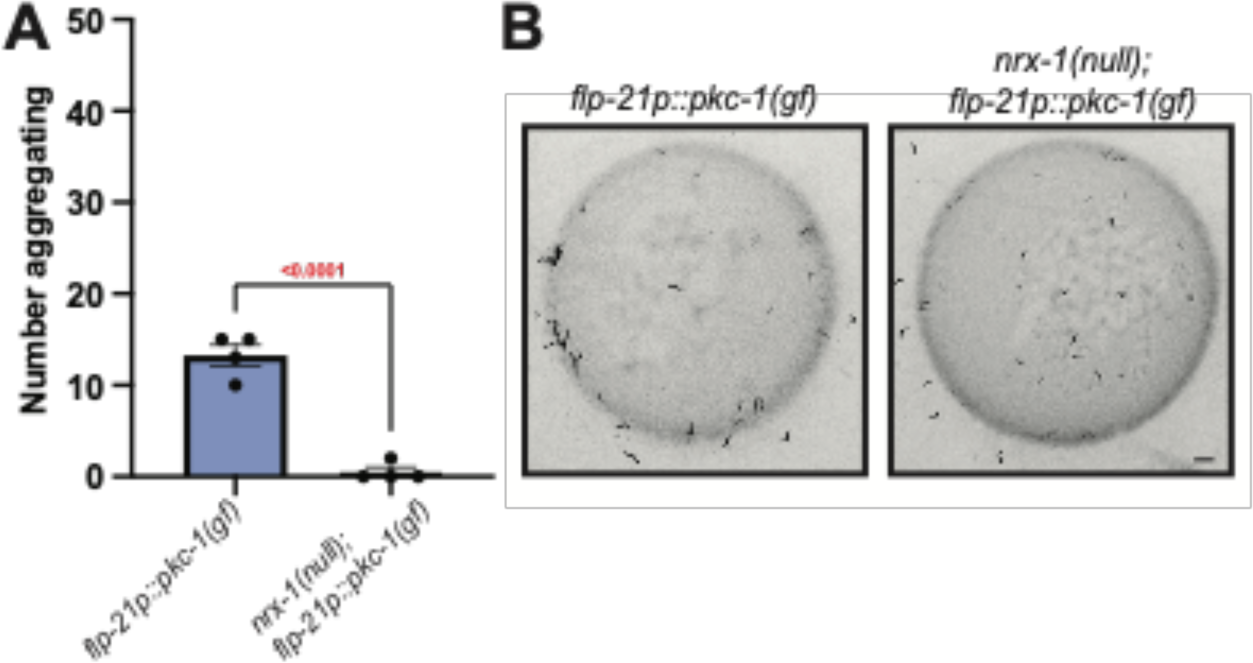
Disruption of *nrx-1* prevents aggregation behavior induced by activation of sensory neurons and RMG interneurons. **A)** Graph showing number of aggregating animals in *flp-21p::pkc-1(gf)* strain compared to *flp-21p::pkc-1(gf);nrx-1(wy778)*. **B)** Representative images of aggregation behavior in *flp-21p::pkc-1(gf)* and *flp-21p::pkc-1(gf);nrx-1(wy778)* animals.

## DISCUSSION

In this study, we identify the mechanisms by which neurexin molecules regulate synapses, neuron signaling, and social feeding behavior. In doing so, we identify multiple signaling pathways that modify the synaptic properties of sensory neurons and tune feeding behavior from social to solitary. We find that the conserved synaptic signaling genes neurexin (*nrx-1*) and neuroligin (*nlg-1*) are necessary for high aggregation behavior. These genes have an additive impact on behavior, implying that they function independently in social feeding behavior. This is surprising as neurexins and neuroligins are thought to be localized to pre- and post-synapses respectively, and canonically bind each other. However, our results are consistent with those observed in *Drosophila* where *dnl2;dnrx ^Δ^*^83^ double mutants show worsened neuromuscular junction morphologic phenotypes and lethality compared to either mutant alone^85^. We suggest that NLG-1 functions broadly in neurons although expression of NLG-1 in neurons was not sufficient to restore aggregation behavior to levels of *npr-1* animals. This may be due to mis-expression in all neurons, levels or timing of expression, or suggest roles for *nlg-1* in non-neuronal cells, aligning with known post-synaptic functions^86^. Despite the ubiquitous expression of *nrx-1*, we localize *nrx-1* function in aggregate feeding to just two pairs of sensory neurons, ASH and ADL, within a well-studied sensory integration circuit. The *C. elegans* neurexin locus encodes multiple isoforms including orthologs of the mammalian *NRXN1* alpha (α) and gamma (γ) isoforms and our analysis identifies a specific role for the alpha (α) isoform at ASH and ADL synapses to affect aggregation behavior. We show that neurexin signaling acts in parallel with glutamate signaling from ASH and ADL neurons to control aggregate feeding behavior. Double mutants of neurexin (*nrx-1*) and the vesicular glutamate transporter (*eat-4*), have an additive effect on social feeding, compared to single mutants in either gene. Using genetic methods, we also identify both excitatory (AMPA-like *glr-1* and *glr-2*) and inhibitory (glycine receptor-like chloride channel *avr-15*) glutamate receptors that contribute to social feeding behavior. We queried the expression of each receptor and find that downstream of ADL and ASH*, glr-1* and *glr-2* are expressed in command interneurons (AVA, AVE, AVD) and AIB, which control backward locomotion and high angle turning, while *avr-15* is expressed in AIA, which inhibits turning^73,87,88^. We suggest that glutamate release from ADL and ASH neurons act on these glutamate receptors to maintain animal position within the social aggregate. Furthermore, these genes (*nrx-1*, *eat-4*, *glr-1*, *glr-2*, and *avr-15*) result in intermediate reductions in aggregation behavior, distinct from many previously reported mechanisms and loci. Whereas loss of sensory transduction channel subunits (*tax-2*, *tax-4*, *osm-9*, and *ocr-2*) and trafficking machinery (*odr-4* and *odr-8*) abolish aggregate feeding, the signaling mechanisms we identify reduce, but do not eliminate, social feeding. These findings imply that there are distinct pathways that tune behavior from solitary to aggregate feeding, as observed across wild isolates^20^.

Gap junctions and neuropeptide signaling are crucial for *C. elegans* aggregate feeding behavior; but roles for chemical synaptic signaling have not been extensively characterized. We find that aggregating animals have both increased numbers of pre-synaptic puncta and faster rates of glutamate release from ASH neurons compared to their solitary counterparts. While ASH glutamate release dynamics in aggregating animals are not impacted by loss of *nrx-1*, we show the number of ASH pre-synapses depends on the α-isoform of NRX-1, highlighting the complex molecular and circuit mechanisms underlying aggregation behavior. We do not observe any changes in ADL pre-synapses in aggregating animals, or in *nrx-1* mutants, and suggest that ADL may act as an amplifier for ASH based on the bi-directional chemical synapses between ADL and ASH. Our finding that *nrx-1* modifies pre-synaptic puncta number in ASH matches the general role for neurexins in the development and maintenance of pre-synaptic structures. While neurexins are broadly implicated in chemical synaptic properties and social behavior, rarely has gene function, and a single isoform (NRX-1(α)), been simultaneously tied to both circuit mechanisms and behavior. Collectively, our studies identify a role for NRX-1(α) in pre-synaptic architecture of specific synapses (from ASH), separately from glutamate release dynamics, in tuning aggregate feeding behavior.

The number of pre-synaptic release sites and the rate of release represent distinct, but related, mechanisms for regulating chemical synaptic signaling. We propose a tuning model in which glutamate signaling from ASH/ADL positively correlates with level of aggregate feeding. High signaling via ASH in social animals can be lowered either via (1) reduction of ASH synaptic puncta or (2) decrease in the rate of glutamate release, which can be further reduced by these two mechanisms acting together. Loss of *nrx-1*, *nlg-1*, *eat-4, glr-1*, *glr-2*, or *avr-15* alone lead to intermediate levels of aggregation behavior, but combination of two pathways produces more solitary-like behavior through distinct circuit functions. We suggest that ASH glutamate signaling acts as a dial for aggregation behavior, with the increased glutamate neurotransmission (via release rate or sites) driving aggregation behavior and vice-versa. An extension of this model is that it is not glutamate signaling, but rather the overall activity level between sensory neurons and RMG interneurons that controls aggregation behavior. This model would explain how multiple sensory neurons (URX, ASK, ADL, ASH), modalities (oxygen, pheromones, aversive stimuli), and signaling components (NPR-1 inhibition, gap junctions, neuropeptides, release sites, exocytosis) function in the same behavior^21–28,30,33^. Lastly, we show that *nrx-1* is needed for aggregation behavior induced by activation of sensory neurons and RMG interneurons and fits a model where *nrx-1* functions to tune aggregation behavior by regulating neurotransmission and/or neuropeptide release. This also confirms the role for *nrx-1* in aggregation behavior independent of manipulations of *npr-1*.

Aggregate feeding involves the interaction of individual *C. elegans* with each other, matching a definition of social behavior^2^. However, since the first publication of aggregate feeding^20^, there has been a general skepticism about whether this behavior is social^89,90^. Studies have shown that oxygen is an important cue in maintaining these aggregates^23–27^, implying that this behavior might be driven by environmental cues. In contrast, other studies showed that pheromones and touch are also important for aggregation behavior ^21^, suggesting a role for inter-individual interactions in this behavior. Moreover, *C. elegans* can participate in other behaviors that are canonically social. While *C. elegans* exist primarily as self-reproducing competent hermaphrodites, male *C. elegans* also exist. These males are attracted to hermaphrodites through pheromone and ascaroside signaling, prompting mate search and mating^91,92^ – clear examples of social behaviors. Additionally, adult hermaphrodites leave the bacterial food lawn in the presence of their larval progeny, likely to increase food availability to their developing offspring^93^. This potential parental response was shown to depend on nematocin, the *C. elegans* version of the “social hormone” oxytocin^93^ and interestingly also *nlg-1*^94^. Despite these examples of social behavior in *C. elegans* and the involvement of both environmental and social cues in aggregate feeding, the social drive to feed in groups remains controversial.

Variants in human neurexins (*NRXN1*) and neuroligins (*NLGN3*) are associated with increased risk for autism (**Supplementary Table 1**), a neurodevelopmental condition characterized by altered social and communication behaviors, repetitive behaviors, and sensory processing/sensitivity^48,49^. Importantly, through our mechanistic exploration of the social feeding circuit and behavior, we uncovered novel roles for additional conserved autism-associated genes, including *GRIA1,2,3(glr-1)*^95–97^, *GRIA2(glr-2)*^95–97^, and *GLRA2,GABRA3(avr-15)*^98–100^ (**Supplementary Table 1)**^101,102^. Rodent models for some of these genes also implicate them in social behaviors^103–106^. The involvement of these multiple conserved autism-associated genes, which affect social behaviors in mice, rats, and humans, may lend support for aggregate feeding as a simple form of social behavior. Variation in these genes in humans include many genetic changes, often in heterozygous state, whereas here, and in other model organisms, the genes are often studied in the homozygous loss of function context. Importantly, the functional study of autism-associated genes we present does not provide a *C. elegans* model of autism or autism behaviors, which are human specific. Rather, we leverage this pioneering genetic organism, its compact nervous system, and the evolutionarily important social feeding behavior to understand the circuit and molecular mechanisms by which behaviors are modified by conserved genes. These detailed mechanistic discoveries provide a framework to explore the molecular functions of autism-associated genes in social behaviors in more complex model systems and have implications for the autism and neurodiverse communities.

Taken together, this work identifies multiple mechanisms that tune feeding behavior between social and solitary states. We define independent genetic pathways involving many conserved autism-associated genes and chemical signaling mechanisms, including glutamate release dynamics and pre-synaptic structural plasticity, that cooperate to determine foraging strategy. Our work suggests conserved roles for autism-associated genes in driving group interactions between animals across species and provides a mechanistic insight into how these genes control neuronal and circuit signaling to modulate behavior. Lastly, our identification of conserved genes with known roles in social behavior suggest a social origin for aggregate feeding in *C. elegans*^107^.

## METHODS

### *C. elegans* strain maintenance

All strains were maintained on Nematode Growth Medium (NGM) plates and seeded with *Escherichia coli* OP50 bacteria as a food source^108^. Strains were maintained on food by chunking and kept at ∼22-23°C. All strains and mutant alleles included are listed in **Supplementary Table 2** by Fig. order. Solitary controls consist of either N2 strain or transgenic strains expressing reporter constructs in the N2 background and/or *him-8(e1489)* mutation indicated in **Supplementary Table 2**, and aggregate feeding controls consist of DA609 with *npr-1(ad609)* or *npr-1(ad609)* with added reporters and/or *him-8(e1489)* mutation as indicated in **Supplementary Table 2**. Presence of the endogenous *unc-119(ed3)* mutant allele, which was used in the generation of TV13570 *(nrx-1(wy778)),* was not confirmed in our strains. The presence of *him-8(e1489)* and *otIs525[lim-6^int4^p::gfp]*, used in genetic crosses or as an anatomical landmark in indicated Fig. s, did not impact solitary or aggregate feeding behavior (**Supplemental Fig. 1A**). All experiments were performed on hermaphrodites, picked during larval stage 4 (L4), and confirmed as day 1 adults.

### Cloning and constructs

All plasmids are listed in **Supplementary Table 3** along with primer sequences for each promoter. All plasmids were made by subcloning promoters or cDNA inserts into plasmids by Epoch Life Science Inc. as described below. Plasmids for *nrx-1(α)* transgenes were generated by subcloning each promoter to replace the *ric-19* promoter in pMPH34 (*ric-19::sfGFP::nrx-1(α)*), which includes a Superfolder GFP tag fused to the N-terminus of the long α isoform of *nrx-1*. Plasmids for *nlg-1* transgenes were generated by subcloning super folder GFP (primers: fwd - CTGCCCAGGATACGATCCATGAGCAAAGGAGAAGAAC ; rev - AGATCCAGATCCGAGCTCTTTGTAGAGCTCATCC) to replace the N-terminal GFP11 fragment tag on *nlg-1* in plasmid pMVC3^109^, then the *ric-19* promoter was subcloned ahead of the artificial intron and start site of the resulting plasmid (primers: fwd - GCGCCTCTAGAGGATCCcattaaagagtgtgctcca ; rev - TTTGGCCAATCCCGGgttcaaagtgaagagc). The plasmid pMPH45 includes the *ric-19* promoter and a superfolder GFP tag fused to the N-terminus of *nlg-1* (*ric-19::sfGFP::nlg-1*), which was subcloned with indicated promoters to replace the *ric-19* promoter. Plasmids for *eat-4* transgenes were generated by subcloning indicated promoters to replace the *sre-1* promoter in pSM plasmid (*sre-1p-eat-4::sl2::gfp*). To generate plasmids for *cla-1* transgenes, promoters indicated were subcloned to replace the *lim-6 ^int^*^4^ promoter in pMPH21 (*lim-6 ^int^*^4^*::gfp::cla-1*)^77^.

### Transgenic animals

All plasmids and co-injection markers are indicated in **Supplemental Table 2** and were injected to generate extrachromosomal arrays at 20 ng/µl^-1^ unless otherwise indicated in **Supplemental Table 2**. For extrachromosomal transgenes, at least 2 independent transgenic lines were generated and analyzed to confirm expression levels and transmittance, after which a single line was selected for comprehensive analysis based on expression levels and moderate to high transmittance^110^.

### Aggregate feeding behavior assay

Standard 6-well plates were filled with 6mL of NGM. 75 μL of OP50 bacteria culture (OD600 = ∼0.7) was added to the center of the well to form a circular food lawn. Plates were left at room temperature to dry. The day after seeding OP50, 60 L4 hermaphrodites of each genotype were moved to a clean plate then 50 animals were transferred to the aggregation behavior assay set-up. If transgenic strains were used, transgene positive animals were identified by the presence of a fluorescent co-injection marker (listed in **Supplemental Table 2**). *C. elegans* were transferred to the center of the food lawn on each well. Experimenter was blinded to all genotypes at the time of loading. 10x Tween was put on the lid of the 6-well plate to prevent condensation from forming. Loaded 6-well plates were placed in the WormWatcher set up developed by Tau Scientifics and the Fang-Yen Lab at the University of Pennsylvania and monitored for at least 15 hours. Images were taken every 5 seconds for 1 minute per hour.

To quantify aggregation behavior, the number of aggregating *C. elegans* were manually counted from blinded images, such that a *C. elegans* in contact with two or more other *C. elegans* was considered aggregating. In cases where the number of aggregating animals could not be clearly counted, the number of single animals were counted and subtracted from 50 to obtain a count of aggregating *C. elegans*. Data shown is from hour 15 after experimental set-up, therefore representing day 1 adult animals.

### Confocal Microscopy

#### Transgenic Expression

For visualization of transgenic constructs, 5% agar was used to create a thin pad on a microscope slide. 5μl of the paralytic, sodium azide was pipetted on the agar pad. Adult animals expressing the co-injection markers were identified on the florescent microscope and moved to the agar pad and a coverslip was placed on top. *C. elegans* were imaged at 63X on a Leica SP8 point scanning Confocal Microscope, with z-stack images taken at 0.6μm spanning expression. Images were processed in Adobe Photoshop to alter orientation and invert color. Fig. s were made in Adobe Illustrator.

#### CLA-1 Puncta Quantification

Relevant mutant strains were crossed with *psrv-3: cla-1*::*sfGFP* or *psra-6::cla-1::sfGFP* in *him-5* background. To visualize CLA-1::GFP puncta in ADL or ASH, microscope slides were prepared as described above. *C. elegans* were again imaged at 63X, with an additional zoom of 2.5 X and a Z-stack size of 0.6μm. Following imaging, Lightning Deconvolution was applied to the images to reduce noise. The number of puncta was examined in FIJI using Particle Analysis. Image stacks were combined to create a single image using a projection of max intensity. Images were auto-thresholded with a minimum of 50 and maximum of 255. Region of interest for particle quantification was restricted to expression of *cla-1::gfp* in the nerve ring and were drawn to exclude any background. If background fluorescence, resulting from the *lin-44::gfp* co-injection marker in these transgenic strains, was too high to distinguish puncta, images were not quantified. Particle analysis was performed with an area cut-off of 0.03μm^2^ to remove small background particles and bare outlines were generated.

#### Fluorescent Recovery After Photobleaching (FRAP)

For FRAP imaging, L4 *C. elegans* were picked 24 hours before imaging to appropriately stage the animals. The next day, no more than three *C. elegans* were placed on each microscope slide and paralyzed with 5mM levamisole. Using the microlab FRAP module on the Leica SP8 Confocal Microscope at 63X with a zoom of 4.5, a 10μm X 10μm bleach area was defined, centered on the brightest part of the neurite. A recording session was set such that 10 frames were taken pre-blech, 10 frames were taken with 50% laser power applied to the sample, and 138 frames were taken post-bleach with an interframe interval of 0.87 seconds for a total post-bleach recording of two minutes. During the two-minute recovery, animals were monitored to ensure they stayed in frame. If drift was seen, minor manual adjustments to the z-plane were made to hold them in position. If drift was significant or if the animal moved, recording was stopped and not included in our analysis.

To quantify the fluorescence recovery, all traces for each genotype were analyzed using the Stowers Institute Jay Plugins in FIJI^111^. Bleach region was set and fluorescence at each frame was plotted. Graphs were then normalized with the maximum fluorescence set at 1 and the minimum set at 0.

### Statistics and reproducibility

All statistical analyses were performed, and all data were plotted using GraphPad Prism 9.

For behavioral experiments, the hour 15 counts of aggregating animals were plotted for each genotype. Each data point represents an individual well of a 6-well plate. At least four replicates were performed on at least three individual days per genotype. Plots include the standard error of mean (SEM). To compare aggregation behavior levels across genotypes, a one-way ANOVA was performed with a Tukey’s Post-Hoc test applied. P-values are plotted on each graph. For graphs in which only two genotypes are shown (**Supplemental Fig. 1, Fig. 7**), a t-test was used.

For CLA-1::GFP puncta quantification, the number of puncta from each individual image was plotted with SEM and compared between genotypes using a one-way ANOVA with Tukey’s post-hoc test. Imaging sessions were performed on at least three separate days.

For FRAP experiments the average normalized fluorescence value was plotted in GraphPad Prism 9 by frame post-bleach for each strain starting at frame 21 (frame 0 post-bleach) and ending at 158 (frame 138 post-bleach) with SEM was shown. Fractional recovery data were linearly fitted as previously reported^83^.To determine whether the slopes of recovery plots differed between genotypes a t-test was applied between each pair. Experiments were performed on at least three different days.

## ACKNOWLEDGEMENTS

The authors thank the Autism Spectrum Program of Excellence and the labs of Colin C. Conine, Chris Fang-Yen, David M. Raizen, John I. Murray, and Meera V. Sundaram for their feedback on this project, and specifically Anthony D. Fouad (Tau Scientific) for technical support. We also thank Theodore G. Drivas, Brandon L. Bastien, Michael Rieger, Kathleen Quach, Marc V. Fuccillo, Aubrey Brumback, Jonathan T. Pierce, and members of the Hart and Chalasani labs for comments on the manuscript. Some strains were provided by the CGC, which is funded by NIH Office of Research Infrastructure Programs (P40 OD010440). This work was supported in part by a Graduate research fellowship from NSF (MHC), NIH 1R01MH096881, Nippert Foundation (SHC), the Autism Spectrum Program of Excellence at the Perelman School of Medicine and NIH 1R35GM146782 (MPH).

## Funding

Graduate research fellowship from NSF (MHC)

NIH 1R01MH096881 (SHC)

Nippert Foundation (SHC)

Autism Spectrum Program of Excellence at the Perelman School of Medicine (MPH)

NIH 1R35GM146782 (MPH).

## Author Contributions

MHC, SHC, and MPH conceived and designed the study and experiments, and MHC conducted all behavioral and microscopy experiments. MPH designed and generated cloning and plasmids and transgenic strains. KCR generated transgenic animals and performed genetic studies. MHC processed, analyzed, and interpreted all data. MHC wrote the manuscript with assistance from MPH and SHC, and all authors reviewed, revised, and approved the manuscript.

## Data and materials availability

All data and materials are available upon request to the corresponding author. All data are available in the main text or the supplementary materials.

---

**Supplementary table 1.**
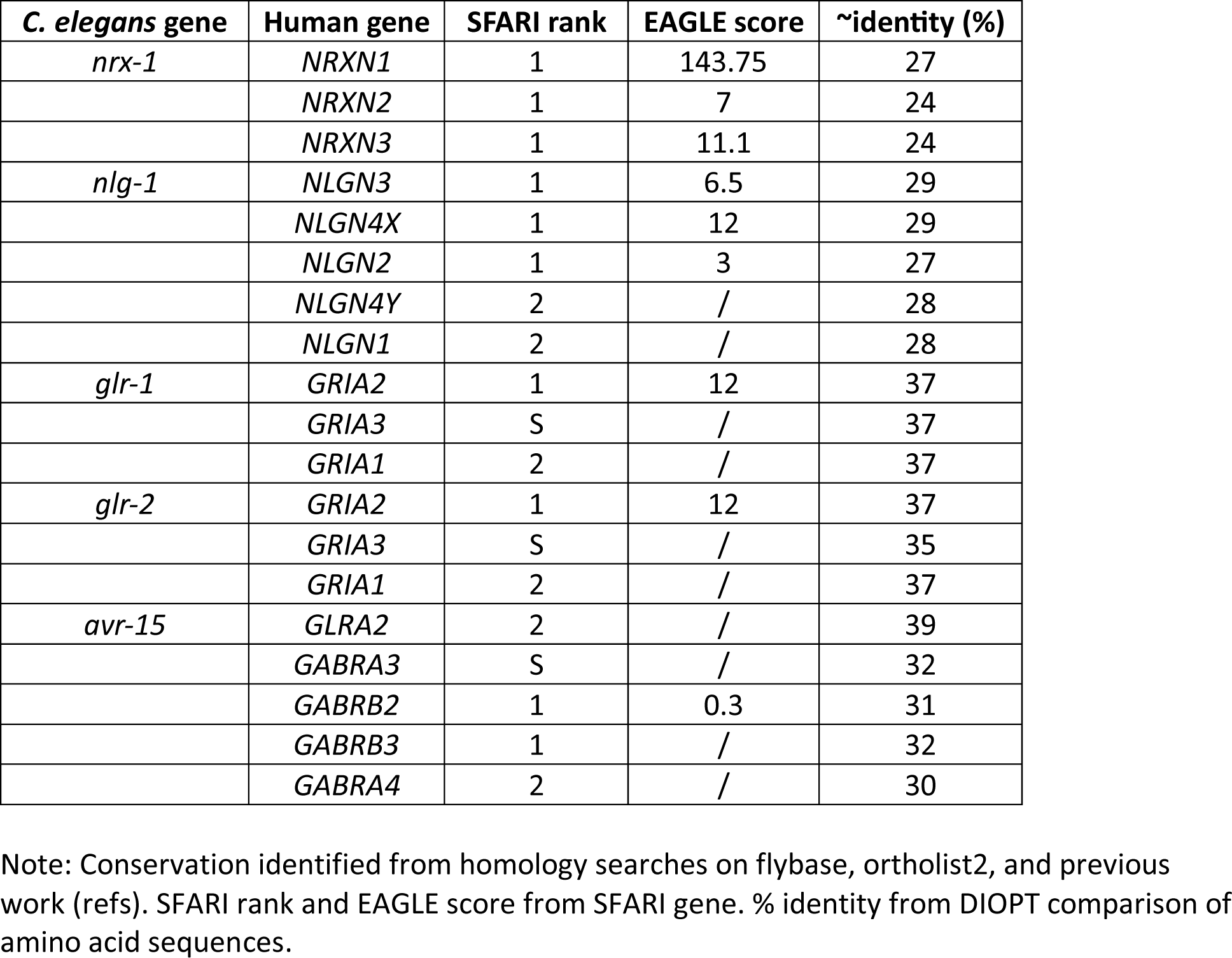
Conservation of *C. elegans* genes with human autism-associated genes.

| *C. elegans* gene | Human gene | SFARI rank | EAGLE score | ~identity (%) |
| --- | --- | --- | --- | --- |
| *nrx-1* | *NRXN1* | 1 | 143.75 | 27 |
|  | *NRXN2* | 1 | 7 | 24 |
|  | *NRXN3* | 1 | 11.1 | 24 |
| *nlg-1* | *NLGN3* | 1 | 6.5 | 29 |
|  | *NLGN4X* | 1 | 12 | 29 |
|  | *NLGN2* | 1 | 3 | 27 |
|  | *NLGN4Y* | 2 | / | 28 |
|  | *NLGN1* | 2 | / | 28 |
| *glr-1* | *GRIA2* | 1 | 12 | 37 |
|  | *GRIA3* | S | / | 37 |
|  | *GRIA1* | 2 | / | 37 |
| *glr-2* | *GRIA2* | 1 | 12 | 37 |
|  | *GRIA3* | S | / | 35 |
|  | *GRIA1* | 2 | / | 37 |
| *avr-15* | *GLRA2* | 2 | / | 39 |
|  | *GABRA3* | S | / | 32 |
|  | *GABRB2* | 1 | 0.3 | 31 |
|  | *GABRB3* | 1 | / | 32 |
|  | *GABRA4* | 2 | / | 30 |

Note: Conservation identified from homology searches on flybase, ortholist2, and previous work (refs). SFARI rank and EAGLE score from SFARI gene. % identity from DIOPT comparison of amino acid sequences.

**Supplementary Table 2.** *C. elegans* strains by Figure.

| Figure | Figure reference | Strain identifier | Genotype | Source | injection concentration* |
| --- | --- | --- | --- | --- | --- |
| 1 | *npr-1* | MPH39 | *him-8(e1489) IV; npr-1(ad609) X; otIs525[lim-6int4::gfp]* | Hart Lab |  |
|  | *npr-1* | DA609 | *npr-1(ad609) X* | CGC |  |
|  | *nrx-1(null); npr-1* | MPH40 | *unc-119(ed3) III; him-8(e1489) IV; nrx-1(wy778[unc-119(+)] V; npr-1(ad609) X; otIs525[lim-6int4::gfp]* | Hart Lab |  |
|  | *nrx-1(null); npr-1* | MPH49 | *unc-119(ed3) III; nrx-1(wy778[unc-119(+)] V; npr-1(ad609) X* | this study |  |
|  | *nrx-1($\alpha$ mut); npr-1* | MPH50 | *him-8(e1489) IV; nrx-1(gk246237) V; npr-1(ad609) X; otIs525[lim-6int4::gfp]* | this study |  |
|  | *nrx-1($\alpha$ del); npr-1* | MPH51 | *nrx-1(nu485) V; npr-1(ad609) X* | this study |  |
|  | solitary controls | OH15098 | *him-8(e1489) IV; otIs525[lim-6int4::gfp]* | Hart & Hobert 2018 |  |
|  | *nrx-1(null)* | TV13570 | *unc-119(ed3) III, nrx-1(wy778[unc-119(+)] V* | CGC |  |
|  | *nrx-1($\alpha$ mut)* | OH15116 | *him-8(e1489) IV; nrx-1(gk246237) V; otIs525[lim-6int4::gfp]* | Hart & Hobert 2018 |  |
|  | *nrx-1($\alpha$ del)* | TV22997 | *nrx-1(nu485) V* | Tong et al. 2017 |  |
| 2 | *npr-1; nrx-1(null); ric-19p::nrx-1($\gamma$)* | MPH52 | *unc-119(ed3) III; nrx-1(wy778[unc-119(+)] V; npr-1(ad609) X; hpmEx3[ric-19p::sfGFP::nrx-1($\gamma$); ttx-3::mCherry]* | this study |  |
|  | *npr-1; nrx-1(null); ric-19p::nrx-1($\alpha$)* | IV870 | *unc-119(ed3) III, him-8(e1489) IV; nrx-1(wy778[unc-119(+)] V; npr-1(ad609) X; otIs525[lim-6int4::gfp]; ueEx601[ric-19p::sfGFP::nrx-1($\alpha$); unc-122p::dsRed]* | this study |  |
|  | *npr-1; nrx-1(null); flp-21p::nrx-1($\alpha$)* | IV874 | *unc-119(ed3) III, him-8(e1489) IV; nrx-1(wy778[unc-119(+)] V; npr-1(ad609) X; otIs525[lim-6int4::gfp]; ueEx605[flp-21p::sfGFP::nrx-1($\alpha$); unc-122::dsRed]* | this study |  |
|  | *npr-1; nrx-1(null); nhr-79p::nrx-1($\alpha$)* | IV878 | *unc-119(ed3) III, him-8(e1489) IV; nrx-1(wy778[unc-119(+)] V; npr-1(ad609) X; otIs525[lim-6int4::gfp]; ueEx609[nhr-79p::sfGFP::nrx-1($\alpha$); unc-122::dsRed]* | this study |  |
|  | *npr-1; nrx-1(null); ric-19p::nrx-1($\alpha$)* | MPH53 | *unc-119(ed3) III; nrx-1(wy778[unc-119(+)] V; npr-1(ad609) X; ueEx611[ric-19p::sfGFP::nrx-1($\alpha$); unc-122p::dsRed]* | this study |  |
|  | *npr-1; nrx-1(null); nhr-79p::nrx-1($\alpha$)* | MPH54 | *unc-119(ed3) III, nrx-1(wy778[unc-119(+)] V; npr-1(ad609) X; ueEx609[nhr-79p::sfGFP::nrx-1($\alpha$); unc-122::dsRed]* | this study |  |
|  | *npr-1; nrx-1(null); srv-3p::nrx-1($\alpha$)* | MPH55 | *unc-119(ed3) III, nrx-1(wy778[unc-119(+)] V; npr-1(ad609) X; hpmEx9[srv-3p::sfGFP::nrx-1($\alpha$); lin-44::gfp]* | this study |  |
| | *npr-1; nrx-1(null); srv-3p::nrx-1($\alpha$); sra-6p::nrx-1($\alpha$)* | MPH56 | *unc-119(ed3) III; nrx-1(wy778[unc-119(+)] V; npr-1(ad609) X; hpmEx9[srv-3p::sfGFP::nrx-1($\alpha$); lin-44::gfp]* | this study | 45 ng/μl |
| | *npr-1; nrx-1(null); sra-* | MPH57 | *unc-119(ed3) III; nrx-1(wy778[unc-119(+)] V; npr-1(ad609) X; him-8(e1489) IV;* | this study | 45 ng/μl |
|  | *6p::nrx-1(α)* |  | *otIs525[lim-6int4::gfp]; hpmEx10[sra-6p::sfGFP::nrx-1(α); unc-122::dsRed]* |  |  |
| 3 | *npr-1; nlg-1* | MPH43 | *him-8(e1489) IV; npr-1(ad609) X; nlg-1(ok259) X; otIs525[lim-6int4::gfp]* | Hart Lab |  |
|  | solitary controls | N2 | Bristol lab control strain | CGC |  |
|  | *nlg-1* | VC228 | *nlg-1(ok259) X* | CGC |  |
|  | *npr-1; nlg-1; ric-19p::nlg-1* | IV930 | *npr-1(ad609) X; nlg-1(ok259) X; ueEx645[ric-19p::sfGFP::nlg-1; lin-44::gfp]* | this study |  |
|  | *npr-1; nlg-1; nhr-79p::nlg-1* | MPH58 | *him-8(e1489)IV; npr-1(ad609) X; nlg-1(ok259) X; otIs525[lim-6int4::gfp]; hpmEx11[nhr-79p::sfGFP::nlg-1; lin-44::gfp]* | this study |  |
|  | *npr-1; nlg-1; nlp-56p::nlg-1* | MPH59 | *him-8(e1489) IV; npr-1(ad609) X; nlg-1(ok259) X; otIs525[lim-6int4::gfp]; hpmEx12[nlp-56p::sfGFP::nlg-1; ttx-3::mCherry]* | this study | 40 ng/μl |
|  | *npr-1; nrx-1; nlg-1* | MPH44 | *unc-119(ed3) III; him-8(e1489) IV; nrx-1(wy778[unc-119(+)] V; npr-1(ad609) nlg-1(ok259) X; otIs525[lim-6int4::gfp]* | this study |  |
| 4 | *npr-1; eat-4* | MPH60 | *eat-4(ky5) III; npr-1(ad609) X; otIs525[lim-6int4::gfp]* | this study |  |
|  | *npr-1; eat-4; nhr-79p::eat-4* | MPH61 | *eat-4(ky5) III; npr-1(ad609) X; otIs525[lim-6int4::gfp]; hpmEx13[nhr-79p::eat-4::SL2::gfp; ttx-3::mCherry]* | this study |  |
|  | *npr-1; eat-4; srv-3p::eat-4* | MPH62 | *eat-4(ky5) III; npr-1(ad609) X; otIs525[lim-6int4::gfp]; hpmEx14[srv-3p::eat-4::SL2::gfp; ttx-3::mCherry]* | this study |  |
|  | *npr-1; eat-4; sra-6p::eat-4* | MPH63 | *eat-4(ky5) III; npr-1(ad609) X; otIs525[lim-6int4::gfp]; hpmEx15[sra-6p::eat-4::SL2::gfp; ttx-3::mCherry]* | this study |  |
|  | *npr-1; nrx-1; eat-4* | MPH64 | *eat-4(ky5) III; unc-119(ed3) III; nrx-1(wy778[unc-119(+)] V; npr-1(ad609) X; otIs525[lim-6int4::gfp]* | this study |  |
|  | *npr-1; nrx-1; eat-4; nhr-79p::eat-4* | MPH65 | *eat-4(ky5) III; unc-119(ed3) III; nrx-1(wy778[unc-119(+)] V; npr-1(ad609) X; otIs525[lim-6int4::gfp]; hpmEx13[nhr-79p::eat-4::SL2::gfp; ttx-3::mCherry]* | this study |  |
|  | *npr-1; nrx-1; eat-4; nhr-79p::nrx-1(α)* | MPH66 | *eat-4(ky5) III; unc-119(ed3) III; nrx-1(wy778[unc-119(+)] V; npr-1(ad609) X; ueEx609[nhr-79p::sfGFP::nrx-1(α); unc-122::dsRed]* | this study |  |
|  | *eat-4* | MT6308 | *eat-4(ky5) III* | CGC |  |
|  | *npr-1; nlg-1; eat-4* | MPH67 | *eat-4(ky5) III; npr-1(ad609) nlg-1(ok259) X; otIs525[lim-6int4::gfp]* | this study |  |
|  | *npr-1; glr-1* | MPH68 | *glr-1(n2461) III; npr-1(ad609) X* | this study |  |
|  | *npr-1; glr-2* | MPH69 | *glr-2(ok2342) III; npr-1(ad609) X* | this study |  |
|  | *npr-1; avr-15* | MPH70 | *avr-15(ad1051) V; npr-1(ad609) X* | this study |  |
|  | *npr-1; nrx-1; glr-1* | MPH71 | *glr-1(n2461) unc-119(ed3) III; nrx-1(wy778[unc-119(+)] V; npr-1(ad609) X; otIs525[lim-6int4::gfp]* | this study |  |
|  | *npr-1; nrx-1; glr-2* | MPH72 | *glr-2(ok2342) unc-119(ed3) III; nrx-1(wy778[unc-119(+)] V; npr-1(ad609) X; otIs525[lim-6int4::gfp]* | this study |  |
|  | *npr-1; nrx-1; avr-15* | MPH73 | *unc-119(ed3) III; nrx-1(wy778[unc-119(+)] avr-15(ad1051) V; npr-1(ad609) X; otIs525[lim-6int4::gfp]* | this study |  |
| 5 | solitary controls | CX16921 | *kyls673[sra-6:eat-4::pHluorin; unc-122:dsRed]* | Bargmann lab |  |
|  | *nrx-1(null)* | MPH74 | *unc-119(ed3) III; nrx-1(wy778[unc-119(+)] V; kyls673[sra-6:eat-4::pHluorin; unc-122:dsRed]* | this study |  |
|  | *npr-1* | MPH75 | *npr-1(ad609) X; kyls673[sra-6:eat-4::pHluorin; unc-122:dsRed]* | this study |  |
|  | *npr-1; nrx-1(null)* | MPH76 | *unc-119(ed3) III; nrx-1(wy778[unc-119(+)] V; npr-1(ad609) X; kyls673[sra-6:eat-4::pHluorin; unc-122:dsRed]* | this study |  |
| 6 | solitary controls | MPH77 | *hpmEx16[srv-3p::gfp::cla-1; lin-44::gfp]* | this study |  |
|  | *nrx-1(null)* | MPH78 | *unc-119(ed3) III; nrx-1(wy778[unc-119(+)] V; hpmEx16[srv-3p::gfp::cla-1; lin-44::gfp]* | this study |  |
|  | *npr-1* | MPH79 | *npr-1(ad609) X; hpmEx16[srv-3p::gfp::cla-1; lin-44::gfp]* | this study |  |
|  | *npr-1; nrx-1(null)* | MPH80 | *unc-119(ed3) III; nrx-1(wy778[unc-119(+)] V; npr-1(ad609) X; hpmEx16[srv-3p::gfp::cla-1; lin-44::gfp]* | this study |  |
|  | solitary controls | MPH81 | *hpmEx17[sra-6:gfp::cla-1; lin-44::gfp]* | this study |  |
|  | *nrx-1(null)* | MPH82 | *unc-119(ed3) III; nrx-1(wy778[unc-119(+)] V; hpmEx17[sra-6:gfp::cla-1; lin-44::gfp]* | this study |  |
|  | *npr-1* | MPH83 | *npr-1(ad609) X; hpmEx17[sra-6:gfp::cla-1; lin-44::gfp]* | this study |  |
|  | *npr-1; nrx-1(null)* | MPH84 | *unc-119(ed3) III; nrx-1(wy778[unc-119(+)] V; npr-* | this study |  |
|  |  |  | *1(ad609) X; hpmEx17[sra-6:gfp::cla-1; lin-44::gfp]* |  |  |
|  | *npr-1; nrx-1($\alpha$ mut)* | MPH85 | *nrx-1(gk246237) V; npr-1(ad609) X; hpmEx17[sra-6:gfp::cla-1; lin-44::gfp]* | this study |  |
| 7 | *flp-21p::pkc-1(gf)* | CX10252 | *kyEx2385[flp-21p::pkc-1(gf)::sl2::gfp; ofm-1::dsred]* | Bargmann lab |  |
|  | *nrx-1(null); flp-21p::pkc-1(gf)* | MPH98 | *unc-119(ed3) III; nrx-1(wy778[unc-119(+)]) V; npr-1(ad609) X; kyEx2385[flp-21p::pkc-1(gf)::sl2::gfp; ofm-1::dsred]* | this study |  |
| Supp 1 | *qglR1* | QG1 | *qglR1 (X, CB4856>N2, npr-1) X* | this study |  |
|  | *qglR1; nrx-1(null)* | MPH86 | *unc-119(ed3) III, him-8(e1489) IV; nrx-1(wy778[unc-119(+)]) V; qglR1 (X, CB4856>N2, npr-1) X; otIs525[lim-6int4::gfp]* | this study |  |
|  | *npr-1; nrx-1(null); osm-6p::nrx-1($\alpha$)* | MPH87 | *unc-119(ed3) III, him-8(e1489) IV; nrx-1(wy778[unc-119(+)]) V; npr-1(ad609) X; otIs525[lim-6int4::gfp]; ueEx603[osm-6p::sfGFP::nrx-1($\alpha$); unc-122::dsred]* | this study |  |
|  | *npr-1; nrx-1(null); sre-1p::nrx-1($\alpha$)* | MPH88 | *unc-119(ed3) III, him-8(e1489) IV; nrx-1(wy778[unc-119(+)]) V; npr-1(ad609) X; otIs525[lim-6int4::gfp]; hpmEx18[sre-1p::sfGFP::nrx-1($\alpha$); unc-122::dsRed]* | this study |  |
|  | *npr-1; nrx-1(null); gcy-36p::nrx-1($\alpha$)* | MPH89 | *unc-119(ed3) III, him-8(e1489) IV; nrx-1(wy778[unc-119(+)]) V; npr-1(ad609) X; hpmEx19[gcy-36p::sfGFP::nrx-1($\alpha$); unc-122::dsRed]* | this study |  |
|  | *npr-1; nrx-1(null); flp-8::nrx-1($\alpha$)* | MPH90 | *unc-119(ed3) III, him-8(e1489) IV; nrx-1(wy778[unc-119(+)]) V; npr-1(ad609) X; hpmEx20[flp-8p::sfGFP::nrx-1($\alpha$); unc-122::dsRed]* | this study |  |
| | *npr-1; nrx-1(null); nlp-56p::nrx-1($\alpha$)* | MPH91 | *unc-119(ed3) III, him-8(e1489) IV; nrx-1(wy778[unc-119(+)]) V; npr-1(ad609) X; otIs525[lim-6int4::gfp]; hpmEx21[nlp-56p::sfGFP::nrx-1($\alpha$); ttx-3::mCherry]* | this study | 40 ng/μl |
|  | *npr-1; nrx-1(null); osm-6p::nrx-1($\alpha$)* | MPH20 | *unc-119(ed3) III; nrx-1(wy778[unc-119(+)]) V; ueEx603[osm-6p::sfGFP::nrx-1($\alpha$); unc-122::dsred]* | this study |  |
|  | *npr-1; nrx-1(null); srv-* | MPH97 | *unc-119(ed3) III, him-8(e1489) IV; nrx-1(wy778[unc-119(+)]) V; npr-1(ad609) X; otIs525[lim-* | this study |  |
|  | *3p::nrx-1(α)* |  | *6int4::gfp*; *hpmEx24[srv-3p::sfGFP::nrx-1(α); lin-44::gfp]* |  |  |
| Supp 2 | *npr-1; nlg-1; sra-6p::nlg-1* | MPH93 | *him-8(e1489) IV; npr-1(ad609) nlg-1(ok259) X; otIs525[lim-6int4::gfp]; hpmEx22[sra-6p::sfGFP::nlg-1; lin-44::gfp]* | this study | 40 ng/μl |
|  | *npr-1; nlg-1; srv-3p::nlg-1* | MPH94 | *him-8(e1489) IV; npr-1(ad609) nlg-1(ok259) X; otIs525[lim-6int4::gfp]; hpmEx23[srv-3p::sfGFP::nlg-1; lin-44::gfp]* | this study | 40 ng/μl |
|  | *npr-1; nlg-1; ins-1p::nlg-1* | MPH92 | *npr-1(ad609) nlg-1(ok259) X; ueEx651[ins-1p::sfGFP::nlg-1; lin-44::gfp]* | this study |  |
|  | *nlg-1; sra-6p::nlg-1* | IV937 | *nlg-1(ok259) X; ueEx651[ins-1p::sfGFP::nlg-1; lin-44::gfp]* | this study |  |
| Supp 3 | *npr-1; nmr-2* | MPH95 | *nmr-2(ok3324) V; npr-1(ad609) X* | this study |  |
|  | *npr-1; mgl-1* | MPH96 | *mgl-1(tm1811) X; npr-1(ad609) X* | this study |  |

\* 20 ng/μl if not noted

**Supplementary Table 3.**
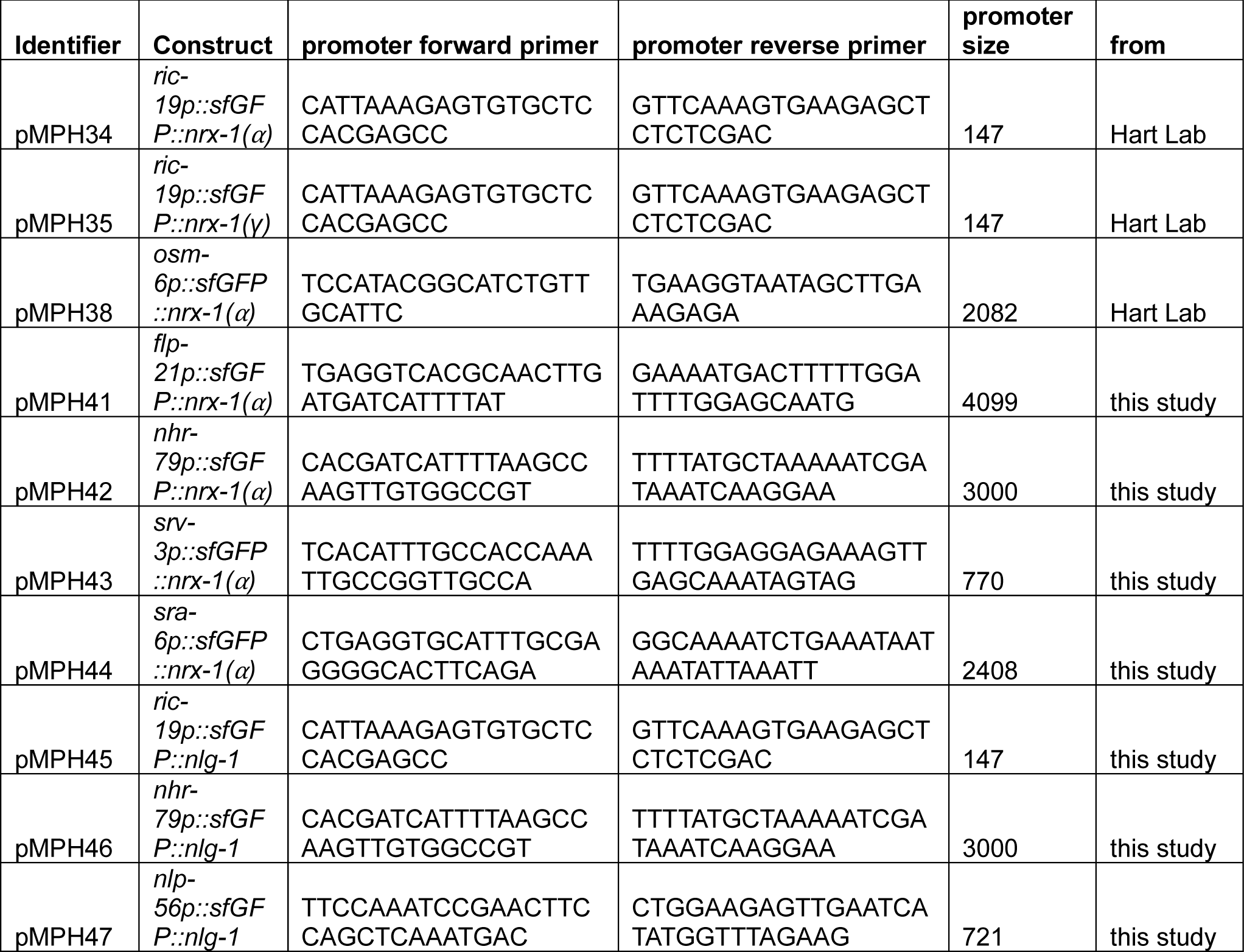
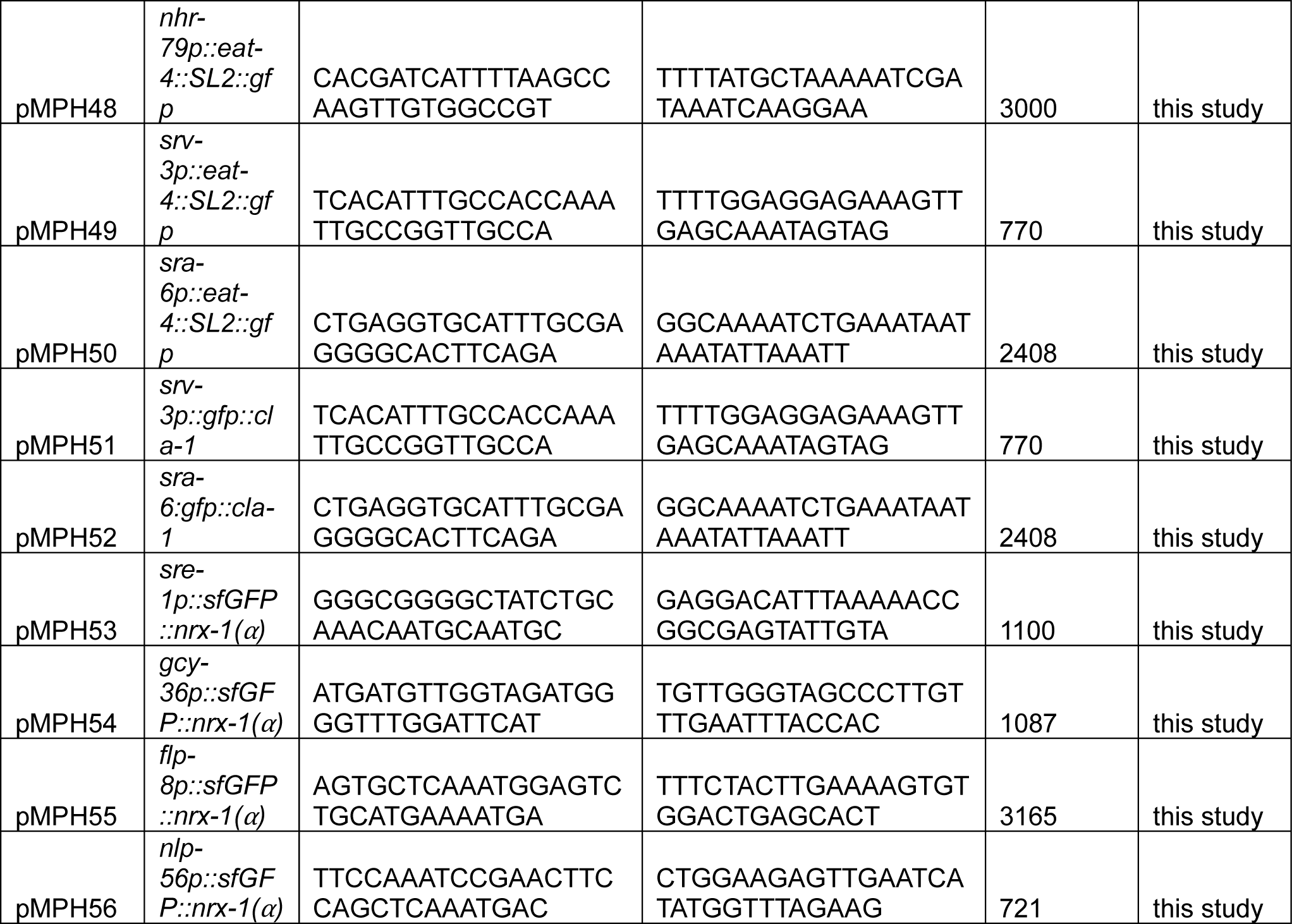
Plasmids and promoters.

**Supplemental Figure 1.**
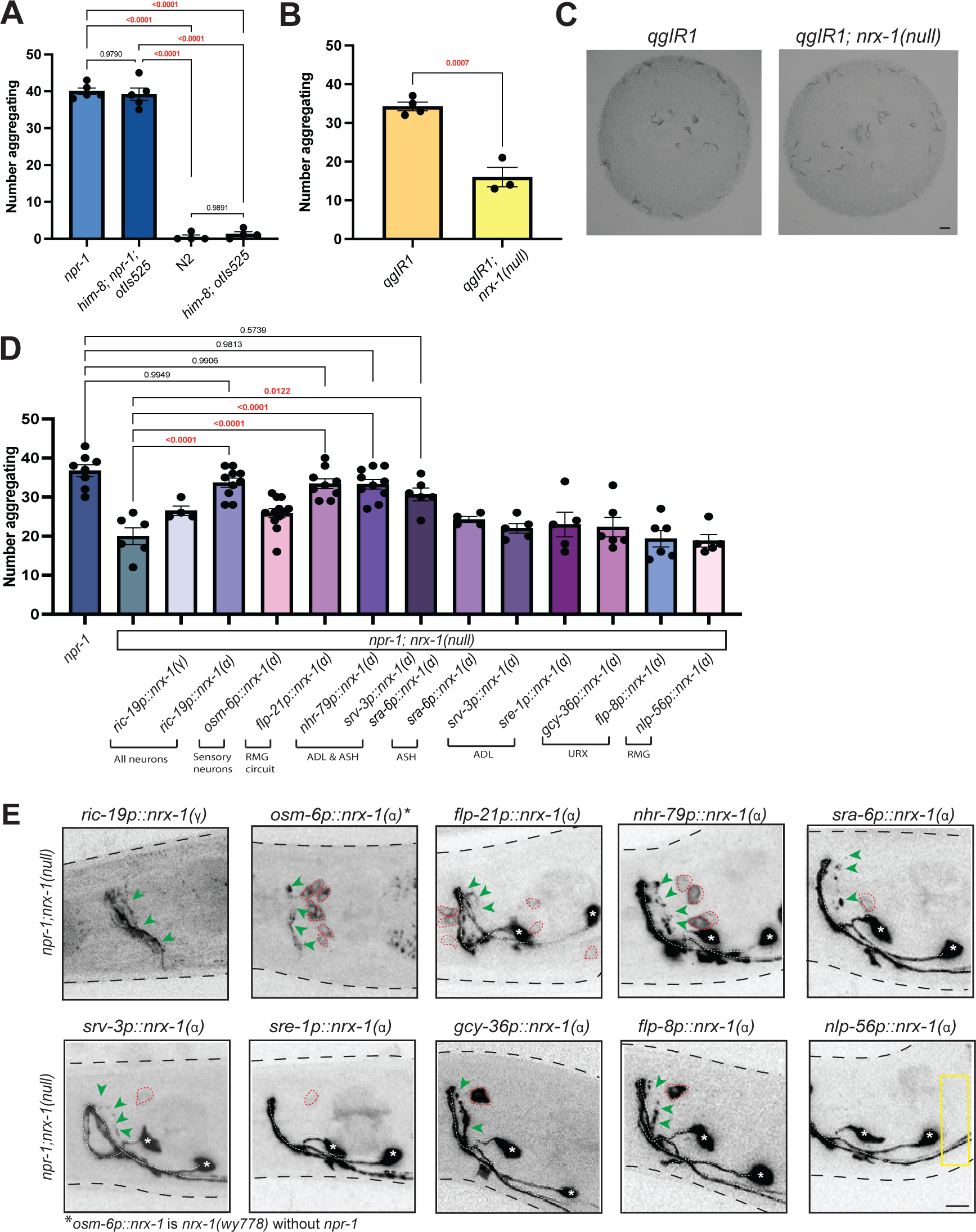
Confirming role for, expression, and localization, of NRX-1 in aggregation behavior. **A)** Graph showing aggregation behavior levels in *npr-1(*ad609) animals compared to *npr-1(ad609); otIs525;him-8* animals and N2 compared to *otIs525;him-8.* Aggregation behavior was not changed by the presence of *otIs525* or *him-8*. Graph showing number of aggregating animals **(B)** and representative images **(C)** of QG1 (*qgIR1*) strain compared to *qgIR1;nrx-1(wy778);otIs525;him-8* mutants (Scale bar = 1mm). **D)** Graph showing number of aggregating animals in *npr-1(ad609), npr-1(ad609);nrx-1(wy778), and npr-1(ad609);nrx-1(wy778)* animals with NRX-1(γ) driven under the *ric-19* promoter and NRX-1(α) driven under promoters indicated. **E)** Expression of NRX-1 tagged with sfGFP driven under various promoters. Green arrows indicate NRX-1 axonal expression. Red dashed lines show cell bodies. White dashed line indicates *lim-6^int^*^4^*::gfp* which drives expression in RIS and AVL axons. White asterisks indicate RIS and AVL cell bodies. Yellow box in *nlp-56p::nrx-1(α)* indicates area where RMG should be located. Expression of *nrx-1* under this promoter is not seen. *osm-6p::nrx-1(α)* imaging performed in *nrx-1(wy778)*(Scale bar = 10μm).

**Supplemental Figure 2.**
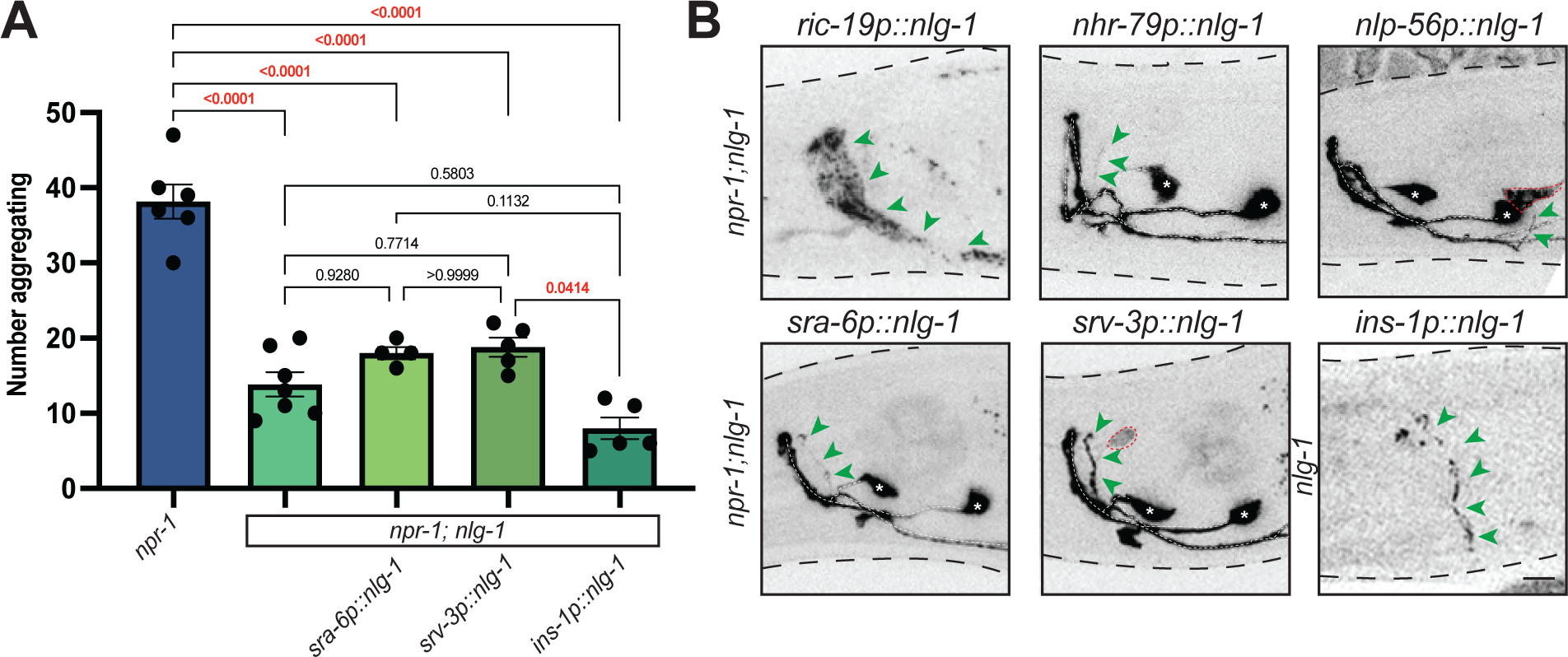
Expression and localization of NLG-1 in aggregation behavior. **A)** Graph showing number of aggregating animals in *npr-1(ad609)*, *npr-1(ad609);nlg-1(ok259)*, and *npr-1(ad609);nlg-1(ok259)* with NLG-1 driven under *sra-6, srv-3*, and *ins-1* promoters. **B)** Expression of NLG-1 tagged with sfGFP driven under various promoters. Green arrows indicate NRX-1 axonal expression. Red dashed lines show cell bodies. White dashed line indicates *lim-6^int^*^4^*::gfp* which drives expression in RIS and AVL axons. White asterisks indicate RIS and AVL cell bodies (Scale bar = 10μm).

**Supplemental Figure 3.**
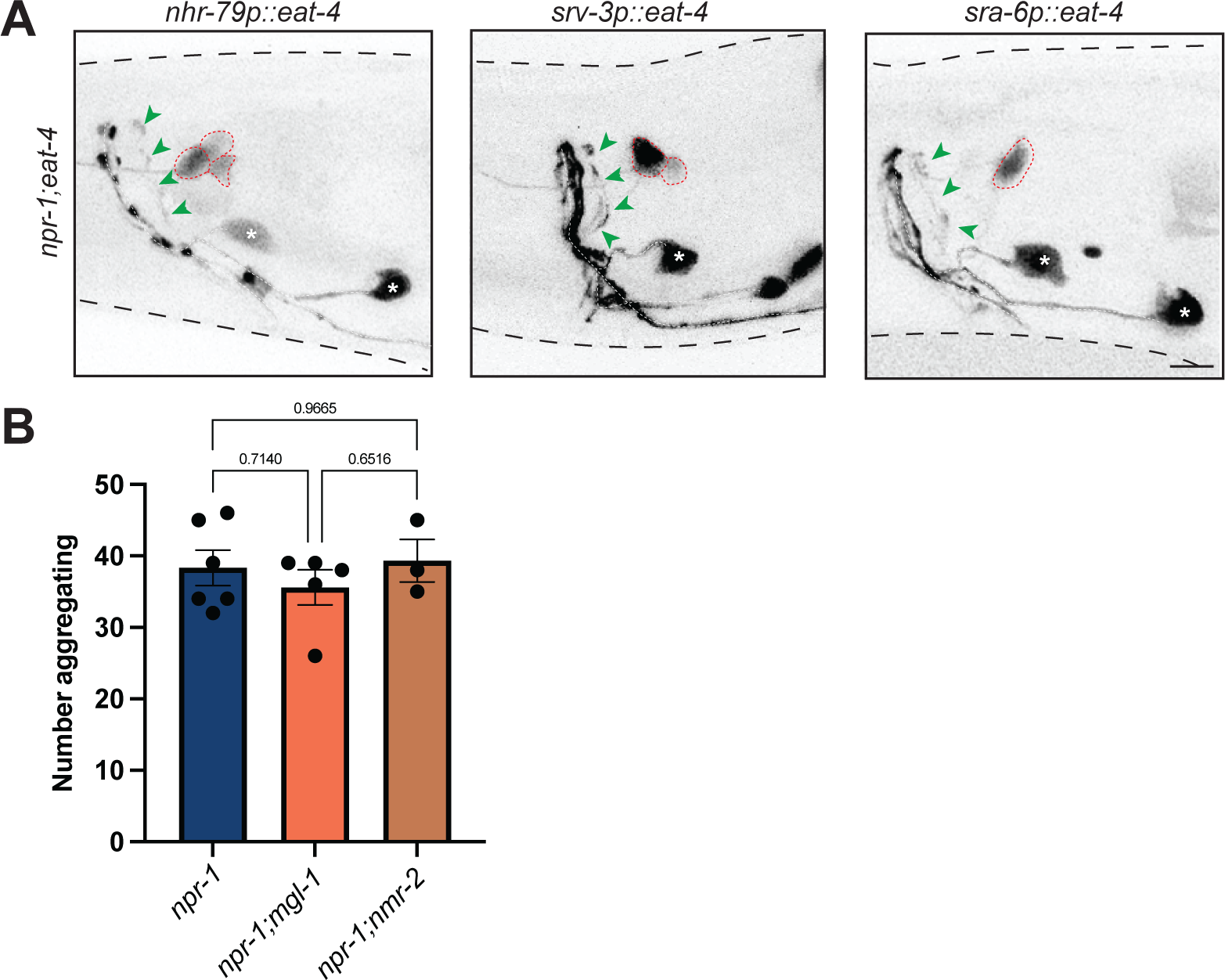
Expression of EAT-4 and analysis of glutamate receptors in aggregation behavior. **A)** Expression of EAT-4 tagged with sfGFP driven under *nhr-79p*, *srv-3p*, and *sra-6p* promoters. Green arrows indicate NRX-1 axonal expression. Red dashed lines show cell bodies. White dashed line indicates *lim-6^int^*^4^*::gfp* which drives expression in RIS and AVL axons. White asterisks indicate RIS and AVL cell bodies (Scale bar = 10μm). **B)** Graph showing number of aggregating animals in *npr-1(ad609),npr-1(ad609);mgl-1(tm1811)* and *npr-1(ad609);nmr-2(ok3324)*.

